# YAP dysregulation triggers hypertrophy by CCN2 secretion and TGFβ uptake in human pluripotent stem cell-derived cardiomyocytes

**DOI:** 10.1101/2024.06.03.597045

**Authors:** Orlando Chirikian, Mohamed A. Faynus, Markus Merk, Zachary Singh, Christopher Muray, Jeffrey Pham, Alex Chialastri, Alison Vander Roest, Alex Goldstein, Trevor Pyle, Kerry V. Lane, Brock Roberts, Jacqueline E. Smith, Ruwanthi N. Gunawardane, Nathan J. Sniadecki, David L. Mack, Jennifer Davis, Daniel Bernstein, Sebastian J. Streichan, Dennis O. Clegg, Siddharth S. Dey, Maxwell Z. Wilson, Beth L. Pruitt

**Affiliations:** Department of Mechanical Engineering, UC Santa Barbara, Santa Barbara, CA, USA; Department of Molecular, Cellular, and Developmental Biology, UC Santa Barbara, CA, USA; Biomolecular Science and Engineering Program, UC Santa Barbara, Santa Barbara, CA, USA; Department of Chemical Engineering, UC Santa Barbara, Santa Barbara, CA, USA; Department of Bioengineering, UC Santa Barbara, Santa Barbara, CA, USA; Department of Physics, UC Santa Barbara, Santa Barbara, CA, USA; Stanford Cardiovascular Institute, Stanford University, CA, USA; Department of Pediatrics (Cardiology), Stanford University School of Medicine, CA, USA; Department of Laboratory Medicine and Pathology, University of Washington, Seattle, WA, USA; Department of Mechanical Engineering, University of Washington College of Engineering, Seattle, WA, USA; Department of Rehabilitation Medicine, University of Washington, Seattle, WA, USA; Institute for Stem Cell & Regenerative Medicine, University of Washington, Seattle, WA 98109, USA; Department of Bioengineering, University of Washington, Seattle, WA 98105, USA; Department of Comparative Medicine, University of Washington, Seattle, WA 98109, USA; Center for Cardiovascular Biology, University of Washington, Seattle, WA 98109, USA; Department of Biomedical Engineering, University of Michigan, Ann Arbor MI, USA; Allen Institute for Cell Science. Seattle, WA, USA

**Keywords:** human induced pluripotent stem cells, cardiovascular disease, cardiomyocytes, hypertrophy, fibrosis, cardiomyopathy, contractile function

## Abstract

Hypertrophy Cardiomyopathy (HCM) is the most prevalent hereditary cardiovascular disease – affecting >1:500 individuals. Advanced forms of HCM clinically present with hypercontractility, hypertrophy and fibrosis. Several single-point mutations in b-myosin heavy chain (MYH7) have been associated with HCM and increased contractility at the organ level. Different MYH7 mutations have resulted in increased, decreased, or unchanged force production at the molecular level. Yet, how these molecular kinetics link to cell and tissue pathogenesis remains unclear. The Hippo Pathway, specifically its effector molecule YAP, has been demonstrated to be reactivated in pathological hypertrophic growth. We hypothesized that changes in force production (intrinsically or extrinsically) directly alter the homeostatic mechano-signaling of the Hippo pathway through changes in stresses on the nucleus. Using human induced pluripotent stem cell-derived cardiomyocytes (hiPSC-CMs), we asked whether homeostatic mechanical signaling through the canonical growth regulator, YAP, is altered 1) by changes in the biomechanics of HCM mutant cardiomyocytes and 2) by alterations in the mechanical environment. We use genetically edited hiPSC-CM with point mutations in MYH7 associated with HCM, and their matched controls, combined with micropatterned traction force microscopy substrates to confirm the hypercontractile phenotype in MYH7 mutants. We next modulate contractility in healthy and disease hiPSC-CMs by treatment with positive and negative inotropic drugs and demonstrate a correlative relationship between contractility and YAP activity. We further demonstrate the activation of YAP in both HCM mutants and healthy hiPSC-CMs treated with contractility modulators is through enhanced nuclear deformation. We conclude that the overactivation of YAP, possibly initiated and driven by hypercontractility, correlates with excessive CCN2 secretion (connective tissue growth factor), enhancing cardiac fibroblast/myofibroblast transition and production of known hypertrophic signaling molecule TGFβ. Our study suggests YAP being an indirect player in the initiation of hypertrophic growth and fibrosis in HCM. Our results provide new insights into HCM progression and bring forth a testbed for therapeutic options in treating HCM.

**Graphical Abstract:** 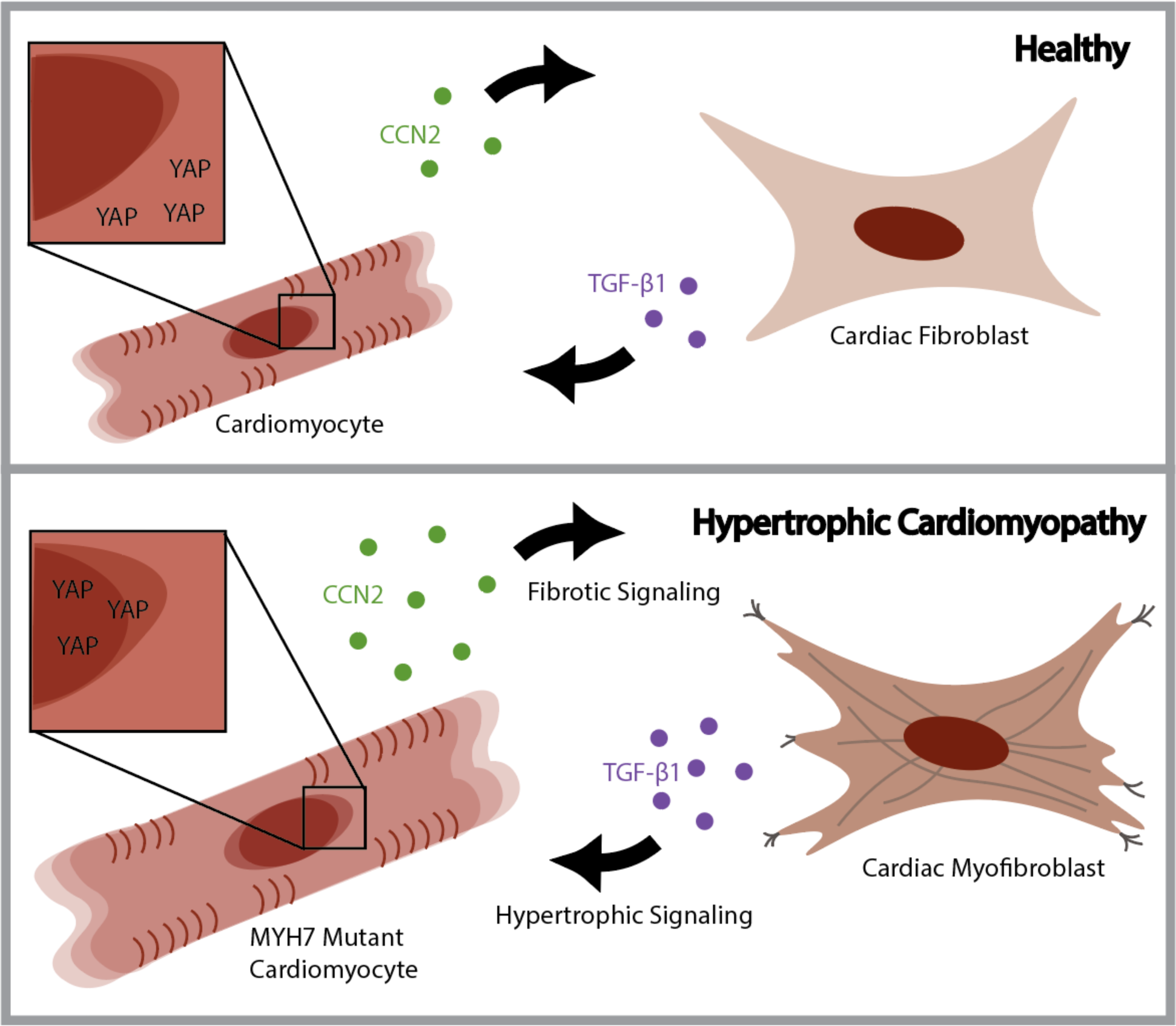

## Introduction

Hypertrophic Cardiomyopathy (HCM) impacts approximately 1 in 500 individuals, making it the most prevalent inherited heart disease^1–4^. The disease manifests as either asymptomatic, permitting patients to reach their full life expectancy, or symptomatic, causing hypertrophied hypercontractile hearts. In symptomatic cases, patients develop left ventricular outflow obstruction and/or systolic dysfunction, which, if left untreated, can lead to heart failure^3–5^. HCM is also one of the leading causes of sudden death due to arrhythmia in adolescents and young adults. The heart’s debilitating hypertrophic growth originates from the hypertrophy of cardiomyocytes rather than their proliferation. In later stages of the disease, myocardial fibrosis emerges as a consequence of the transformation of cardiac fibroblasts into myofibroblasts^6^. Activated myofibroblasts proliferate extensively, accompanied by excessive deposition of extracellular matrix (ECM) proteins, directly contributing to an overall stiffening of the myocardium^7^. While specific mutations have been associated with these characteristic HCM phenotypes, how they interact with hypertrophic and fibrotic signaling pathways remains unknown. The majority of prevalent and well-studied genetic mutations, accounting for around 70% of HCM cases, reside in genes encoding β-myosin heavy chain (MYH7) and myosin-binding protein C (MYBPC3), which both regulate actin-myosin association within the sarcomere^8–10^. Other HCM mutations are situated in alternative sarcomere proteins, suggesting the importance of contractility in driving the phenotypes of the disease. Although clinical interventions capable of slowing or halting the disease’s progression are limited, negative inotropic drugs such as verapamil (a calcium channel blocker)^11–14^ and Mavacamten (a myosin inhibitor)^15,16^ exhibit promising results in diminishing left ventricular hypertrophy. Altogether, the evidence links hypercontractility with cardiomyocyte hypertrophy and fibrosis, leading to a stiffer mechanical environment.

Cardiomyocytes actively experience both inter- and intracellular forces with each contraction^17,18^. These mechanical forces can be exacerbated by cardiac injury or hereditary diseases like HCM^19^. The ability of cells to perceive and react to their environment, known as mechanotransduction, orchestrates the regulation of the epigenome and transcriptome, driving cellular fate^20–22^. The numerous mechanotransducers within cells, including integrins, cadherins, and cytoskeletal networks, directly transmit mechanical signals (forces) to the nucleus via cytoskeletal-LINC protein complexes^23^. As a result, these inter- and intracellular forces can induce alterations in chromatin configuration and accessibility, mediate the transport of transcription factors or transcriptional co-activators from their dormant to active state, resulting in drastic changes to the transcriptome^24–26^. Here, we sought to answer two questions: (1) How does enhanced contractility alter the transcriptomic behavior of cardiomyocytes, and (2) to what extent do these alterations contribute to the phenotypic manifestations of hypertrophic cardiomyopathy?

Given the relationship between altered mechanics and organ growth, the Hippo pathway emerges as a prime candidate regulating the development of HCM. The Hippo pathway, an evolutionarily conserved signaling cascade, governs diverse biological processes including development, cell growth, cell migration, organ size regulation, and tissue regeneration^27–29^. Yes Associated Protein (YAP), a pivotal effector molecule within the Hippo pathway, has been substantiated by multiple research groups as being altered in both human HCM heart tissue samples and murine transverse aortic constriction (TAC) models^30,31^. These studies indicate reduced levels of phosphorylated YAP (i.e. activated YAP with increased nuclear localization) and elevated levels of the well-established YAP downstream transcriptional target, Cellular Communication Network Factor 2 (CCN2)^31^. A recent study leveraging a multi-omic approach to characterize pediatric human patient tissues validates YAP alterations in various congenital heart diseases, reiterating its perturbation in HCM^30^. Additionally, prior investigations describe the activation of cardiac fibroblasts (myofibroblasts), suggesting enhanced pro-fibrotic and pro-hypertrophic Transforming Growth Factor-β (TGFβ) signaling^30,32–34^. However, the defined role of YAP in the development of the HCM phenotypes: hypertrophic enlargement of the cardiomyocytes and of fibrosis remains unclear.

The prevailing influence of mechanical stimuli in the dysregulation of YAP has been associated with several diseases and disease phenotypes, notably those involving proliferation, hypertrophy, regeneration, and migration, such as cancer and various tissue fibrosis^35–38^. This consequently raises our final question: (3) How does a cardiomyocyte modulate YAP activity in a mechanically dynamic environment? Using human induced pluripotent stem cells (hiPSCs) and CRISPR technology, we have successfully recapitulated a hypercontractile phenotype for four HCM mutations in MYH7. Our results suggest a link between YAP and the induction of CCN2 secretion by hiPSC-CMs. CCN2 triggers the activation of cardiac fibroblasts into myofibroblasts, and subsequently the downstream secretion by fibroblasts of the hypertrophic signaling molecule TGFβ1. This feedforward loop exacerbates several features of HCM.

## Results

Despite some limitations due to immaturity, hiPSC derived cardiomyocytes (hiPSC-CMs) have been effectively utilized to model a range of cardiovascular diseases^39–44^. While characterizing the effects of sarcomeric mutations in the manifestation of the hypertrophic phenotype when modeling HCM, researchers have reported mixed outcomes^45^. In this context, we elucidate an explanation for these inconsistencies by showcasing the significance of paracrine signaling from cardiac fibroblasts in the hypertrophic signaling process. We selected three MYH7 mutations having clinical phenotypic occurrence with pronounced pathologies (D239N, H251N, R663H) and one with mixed clinical phenotypes (G256E)^46–50^. Employing a directed cardiac differentiation protocol that modulates the WNT pathway, we effectively differentiated all four mutant lines and respective controls into cardiomyocytes, as evidenced by the expression of the cardiac-specific marker αActinin (Figure 1A)^51^.

**Figure 1.**
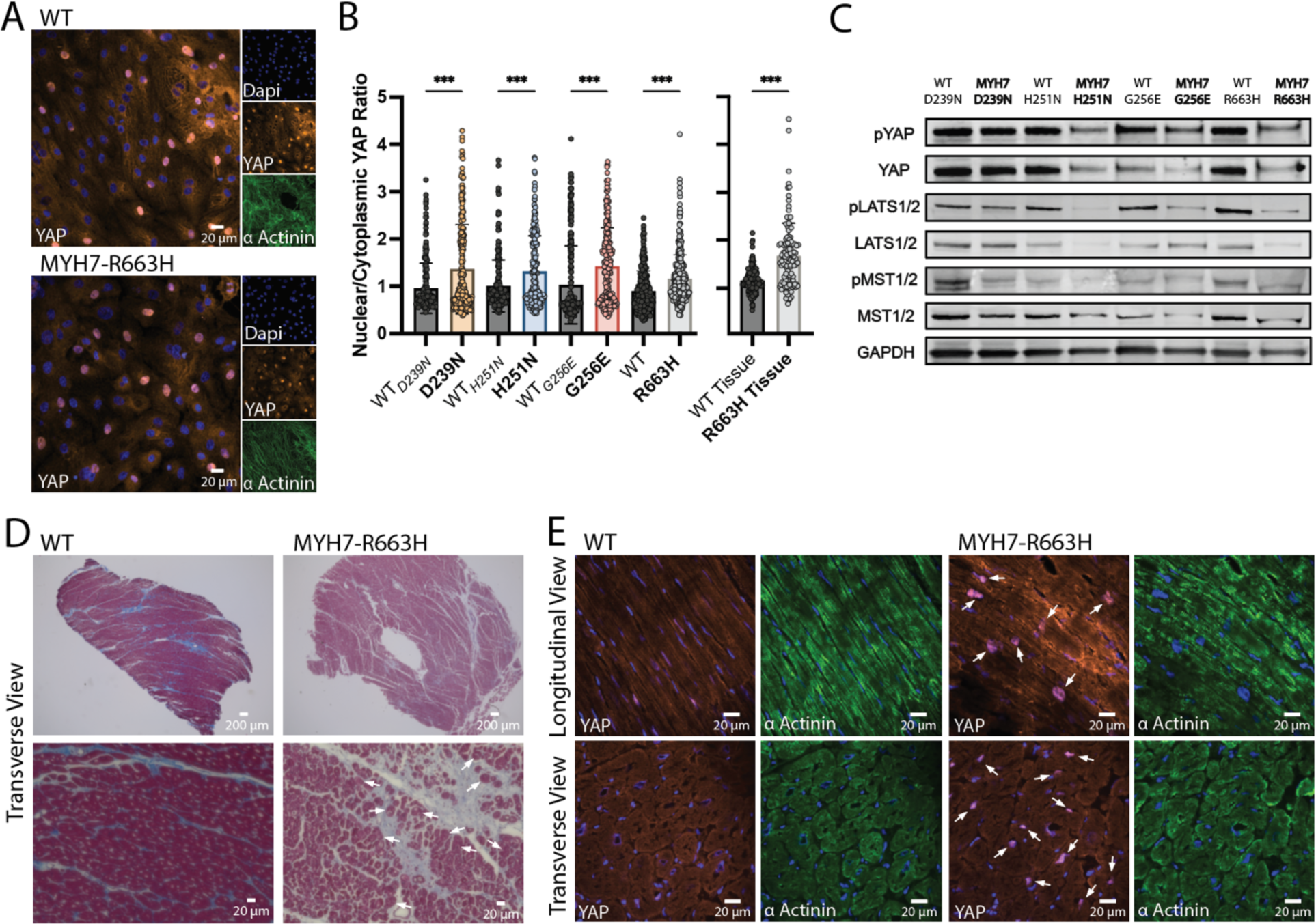
YAP activity is enhanced in hiPSC-CM and human tissue with HCM associated MYH7 mutations. (A) Representative image of fixed and labeled D60 hiPSC-CMs for YAP (red), Alpha Actinin (green) and nuclei (blue) examined by immunocytochemistry. (B) Quantification of nuclear to cytoplasmic YAP expression (n= 200 cells for each cell line, from 3 different differentiations). Data are presented as mean ± STDEV. Statistical test: Mann-Whitney. (C) Representative Western blots probing key Hippo pathway mediates: YAP, pYAP, LATS1/2, pLATS1/2, MST and pMST. GAPDH was utilized as a loading control. (D) Trichrome stained healthy (WT) and diseased (MYH7-R663H) human cardiac tissue. White arrows indicate hypertrophic cardiomyocytes. (E) Fixed and sectioned healthy (WT) and diseased (MYH7-R663H) human tissue labeled for YAP (red), Alpha Actinin (green) and nuclei (blue) evaluated by immunohistochemistry. White arrows indicate nuclear localized YAP. P<0.05 is designated with (*), P<0.005 is designated with (**), P<0.0005 or smaller is designated with (***).

### hiPSC-CMs with HCM-associated mutations have elevated YAP activity

The role of YAP in hypertrophic cardiomyopathy remains uncertain: some propose that increased nuclear YAP in cardiomyocytes initiates cellular hypertrophy, while others suggest a proliferative response^52,53^. To investigate this dichotomy, we examined hiPSC-CMs with HCM-associated mutations for changes in YAP localization using immunocytochemistry. We fixed and stained hiPSC-CMs for YAP on D60 to ensure that maturity would not confound our findings (given YAP’s pivotal role in development)^54^. Intriguingly, the data suggest elevated YAP activity in all four HCM mutant lines compared to their respective controls, inferred from YAP’s nuclear localization (Figure 1A-B).

We next probed the S127-phosphorylated variant of YAP (denoted as pYAP) by Western blot and observed reduced phosphorylated YAP protein levels across all four HCM mutants, consistent with heightened nuclear localization of YAP (Figure 1C)^55^. To further verify YAP’s activity we surveyed the phosphorylated variants of upstream core kinases MST1/2 and LATS1/2. Both MST1/2 and LATS1/2 directly influence YAP localization. Reduced phosphorylation levels of MST1/2 and LATS1/2 in all four HCM mutants (Figure 1C) is also consistent with enhanced nuclear localization of YAP in D60 HCM mutant hiPSC-CMs. Since YAP is alternatively spliced to form multiple isoforms and previous studies have indicated an isoform specific role of YAP, we explored whether MYH7 mutant hiPSC-CMs selectively regulated specific YAP isoforms^56^. RNA purified from WT and HCM mutant D60 hiPSC-CMs was reverse transcribed into cDNA and run separately on a 3% agarose gel to discern distinct band separation characteristic of YAP isoforms, as previously illustrated^56^.

Notably, no discernible changes emerged in band pattern or intensity between WT and HCM hiPSC-CMs, suggesting that isoform expression remains unaltered in the presence of MYH7 HCM-associated mutations (Supplement 1A).

### Human tissue with HCM-associated mutation R663H substantiates enhanced YAP activity in hiPSC model

To validate these findings beyond the hiPSC-CM model system, we examined healthy (WT) and clinically diagnosed HCM human cardiac tissue (MYH7-R663H) for YAP activity. Previous research has highlighted altered Hippo activity within cardiac tissue as a whole, but its specific impact on cardiomyocytes remains unknown. Trichrome staining of MYH7-R663H mutant tissue revealed both cardiomyocyte hypertrophy and increased collagen deposition compared to control tissue, indicative of elevated fibrosis—both characteristic features of advanced HCM (Figure 1D). Immunohistochemistry verified increased YAP nuclear localization, specifically in cardiomyocytes, within MYH7-R663H mutant tissue in contrast to healthy WT tissue. These outcomes paralleled those observed in the hiPSC-CM with the MYH7-R663H mutation (Figure 1B&E, Supplement 1B).

### hiPSC-CMs with MYH7 HCM-associated mutations exhibit a hypercontractile phenotype

One prominent characteristic of HCM is the hypercontractile phenotype displayed by cardiomyocytes. Employing Traction Force Microscopy (TFM), we measured contractile force generation of single patterned hiPSC-CMs on physiological stiffness (10 kPa) polyacrylamide hydrogels (Supplement 2A)^57,58^. Isolated D28 hiPSC-CMs were cultured on hydrogels for 2 days prior to traction force measurements (performed on D30, standard maturity labeled by enhanced expression of MYH7 and TNNI3 and depletion of proliferative mark EdU^59^). The TFM platform enabled the assessment of traction forces for 53-112 cardiomyocytes from three differentiation batches per cell line (four mutant and four respective control lines)^57^.

Peak Force (N), derived from fluorescent bead displacement on hydrogels, displayed a significant increase of at least 2-fold in all four MYH7 mutant cell lines compared to their respective controls (mean fold change: D239N= 2.00, G256E= 3.75, H251N= 4.77, R663H= 4.31) (Figure 2 A-C). Three out of the four MYH7 mutant lines also exhibited significantly enhanced contraction velocity and relaxation velocity (H251N, G256E, and R663H, Supplement 2B-C).

**Figure 2.**
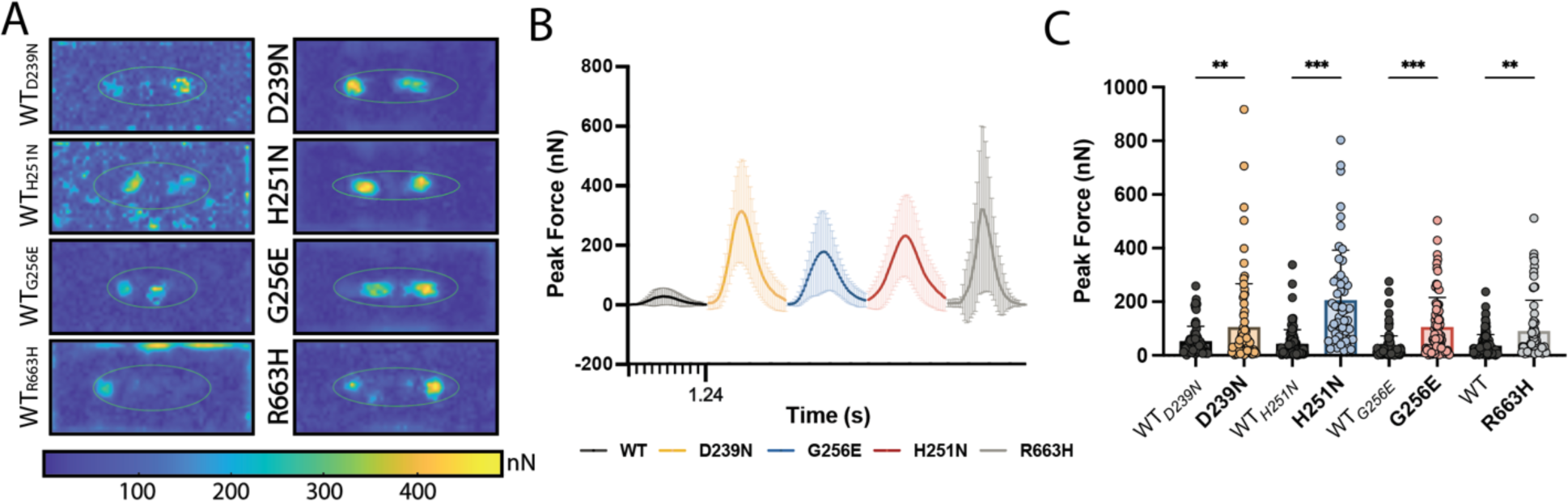
hiPSC-CMs harboring HCM associated MYH7 heterozygous mutations are hypercontractile. (A) Representative Peak Traction Force heatmap plots for each MYH7 mutation and its paired isogenic control. (B) Representative average force trace for each MYH7 mutation depicting the heterogeneity in trace shape for a single differentiation (n=20, average force trace from 5 contraction cycles). Error bars represent STDEV. (C) Summarized quantification of Peak Total Force for each MYH7 mutation D239N (n=77 cells), H251N (n= 53 cells), G256E (n= 82 cells), R663H (n= 63 cells) and its paired isogenic control WT_D239N_ (n= 70 cells), WT_H251N_ (n= 102 cells), WT_G256E_ (n= 108 cells), WT_R663H_ (n= 112 cells). Data are presented as mean ± STDEV. Statistical test: Kruskal-Wallis between mutant and paired isogenic control. P<0.05 is designated with (*), P<0.005 is designated with (**). P<0.0005 or smaller is designated with (***).

### Increasing Contractility drives YAP Nuclear Localization

Prior research has made notable strides in understanding how MYH7 point mutations associated with HCM alter the kinetics of the contractile machinery within cardiomyocytes^8,9,14,60–62^. Our focus revolves around understanding how these modified mechanics influence the state of a cardiomyocyte, particularly in terms of activating or deactivating YAP. We increased the peak force of healthy (WT) hiPSC-CMs by exposing them to substrates mimicking moderate and severe fibrosis (35kPa and 100 kPa); as expected, we also observed increased nuclear YAP (Supplement 2D-E)^63^. We further isolated force production from substrate stiffness by utilizing positive and negative inotropes: levosimendan and verapamil, respectively (Supplement 2F-G)^64,65^. Treatment of WT D60 hiPSC-CMs with force-reducing verapamil (100 nM) for 24 hours led to unchanged YAP nuclear localization (Figure 3A). However, treatment of WT D60 hiPSC-CMs with force-enhancing levosimendan (300 nM) for 24 hours led to significantly increased YAP nuclear localization (Figure 3A). In contrast, when verapamil and levosimendan were administered to D60 MYH7-R663H hiPSC-CMs for 24 hours, a significant reduction in the nuclear localization of YAP was observed following the force-reducing verapamil treatment (Figure 3B). Interestingly, a substantial decrease in the YAP nuclear localization was detected in D60 MYH7-R663H hiPSC-CMs subjected to the force-enhancing levosimendan treatment (Figure 3B). This decrease could be due to the drug’s upstream calcium-sensitizing effect on contraction dynamics, disrupting the mutation-induced alteration in contractility^64^.

**Figure 3.**
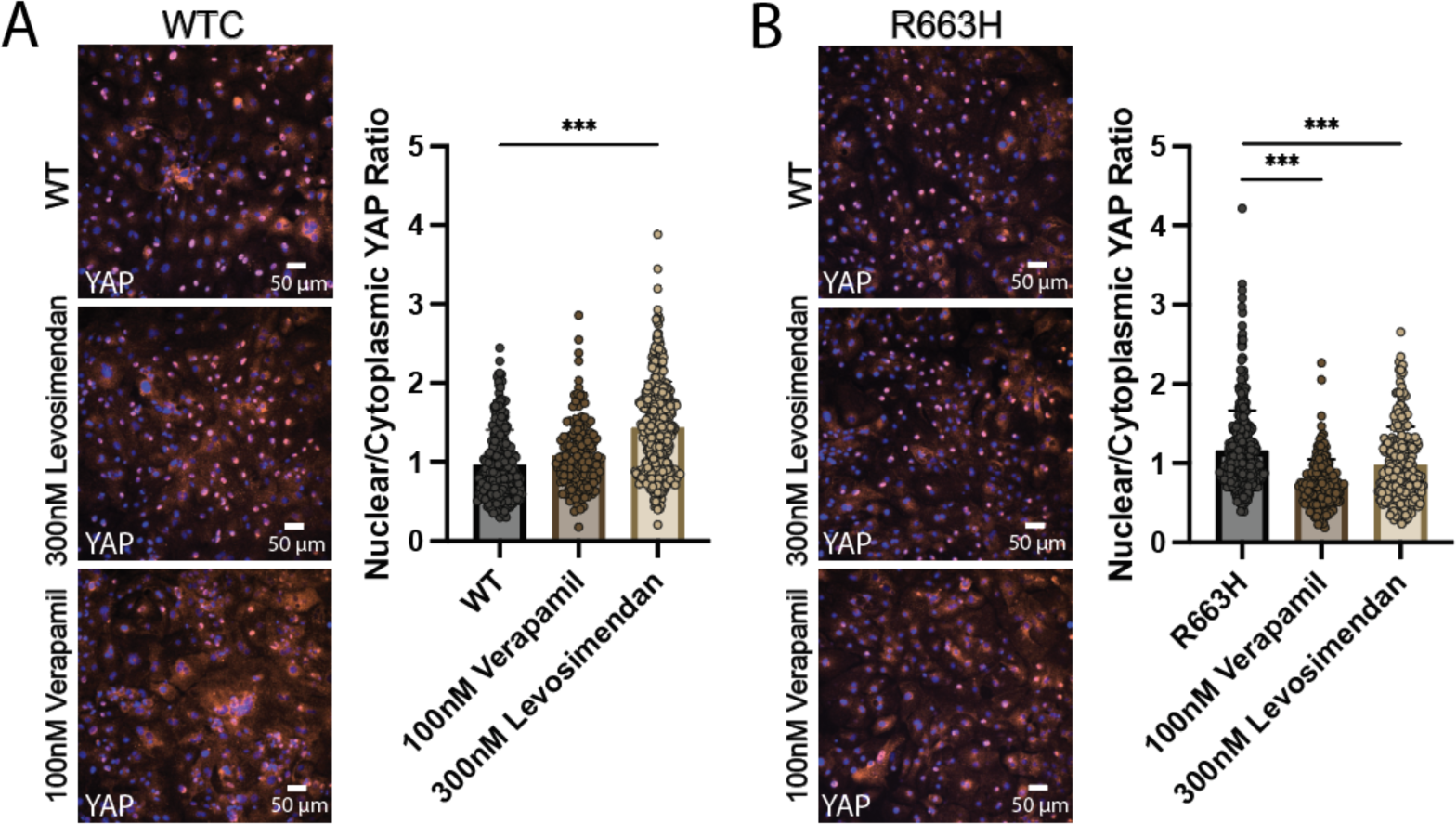
YAP nuclear localization is increased in response to contractility modifying inotropic drugs. (A) Representative images of fixed and labeled WT WTC D60 hiPSC-CMs for YAP (red) and nuclei (blue) treated with either negative or positive inotropic drugs verapamil (100 nM) or levosimendan (300 nM). Quantification of YAP activity from drug experiments inferred by Nuclear to Cytoplasmic intensity Ratio (n= 200 cells for each treatment, from 2 different differentiations). Data are presented as mean ± STDEV. Statistical tested used: one-way ANOVA. (B) Representative image of fixed and labeled disease MYH7-R663H D60 hiPSC-CMs for YAP (red) and nuclei (blue) treated with negative or positive inotropic drugs verapamil or levosimendan. Quantification of YAP activity inferred by Nuclear to Cytoplasmic intensity Ratio (n= 200 cells for each treatment, from 2 different differentiations). Data are presented as mean ± STDEV. Statistical tested used: one-way ANOVA. P<0.05 is designated with (*), P<0.005 is designated with (**). P<0.0005 or smaller is designated with (***).

Together, our data suggest that contractility contributes to the enhanced nuclear localization of YAP. Subsequently, we asked if the mechanical strain from enhanced contraction alone drives the nuclear localization of YAP. To test this hypothesis, we employed a previously established uniaxial stretch device in conjunction with the contraction-inhibiting small molecule blebbistatin, thus enabling the separation of the mechanical strain produced by cardiomyocyte contraction from the intrinsic contraction process (Supplement 3A)^66^. Applying uniaxial static and cyclic (1hz) stretch in alignment with contracting WT hiPSC-CMs substantially heightened the nuclear localization of YAP (Supplement 3B-C). However, stretched and blebbistatin treated WT cardiomyocytes (non-contracting) exhibited significantly reduced nuclear YAP (Supplement 3B-C). These findings suggest that YAP may be driven by the mechanical strain produced by cardiomyocyte contraction. We also tested uniaxial static and cyclic (1hz) stretch perpendicular to WT hiPSC-CMs alignment and observed a significant increase in the nuclear localization of YAP, albeit less than that observed with parallel stretch (Supplement 3D-E).

### Nuclear Deformation is increased in MYH7 Mutant hiPSC-CMs and correlates with YAP nuclear localization

YAP can be activated both by upstream mechanical and/or biochemical signals. Multiple research groups have demonstrated that mechanical stimuli that create observable nuclear deformation exert a more pronounced influence, promoting YAP nuclear localization irrespective of biochemical signals or kinase activity^24,67–70^. Thus, we asked whether the observed endogenous nuclear YAP in MYH7 mutant hiPSC-CMs was accompanied by nuclear deformation changes in relation to enhanced contractility in HCM mutants. Nuclear deformation of contracting hiPSC-CMs was apparent by eye. We quantitatively assessed nuclear aspect ratio change using Hoechst DNA stain and high frame rate video microscopy. We recorded videos of D60 hiPSC-CMs paced at 1 Hz and analyzed them using a custom Fiji script (see Supplement Information). Raw aspect ratio traces shown in Figure 4A were then scrutinized for changes in peak nuclear deformation (amplitude, A) and time under deformation (TD75). Time under deformation was determined by measuring the width (s) of the curve at 75% of the height (from the top, TD75). hiPSC-CMs carrying HCM mutations displayed elevated amplitude and TD75 (Figure 4A-B). Furthermore, both the rise and decay times of deformation were larger (Supplement 4A-B). We then asked whether the increased nuclear deformation drove changes in structural proteins of the nuclear envelope. We assessed gene targets Emerin, Plectin, and Lamin A/C—associated with enhanced nuclear strain— through qPCR (Supplement 4C)^71^. The results suggest an upregulation of Emerin and Plectin in select HCM mutants, along with an upregulation of Lamin A/C in all four mutants. The data suggest a compensatory response by mutant hiPSC-CMs to increase nuclear structural integrity by increasing Lamin A/C transcript levels.

**Figure 4.**
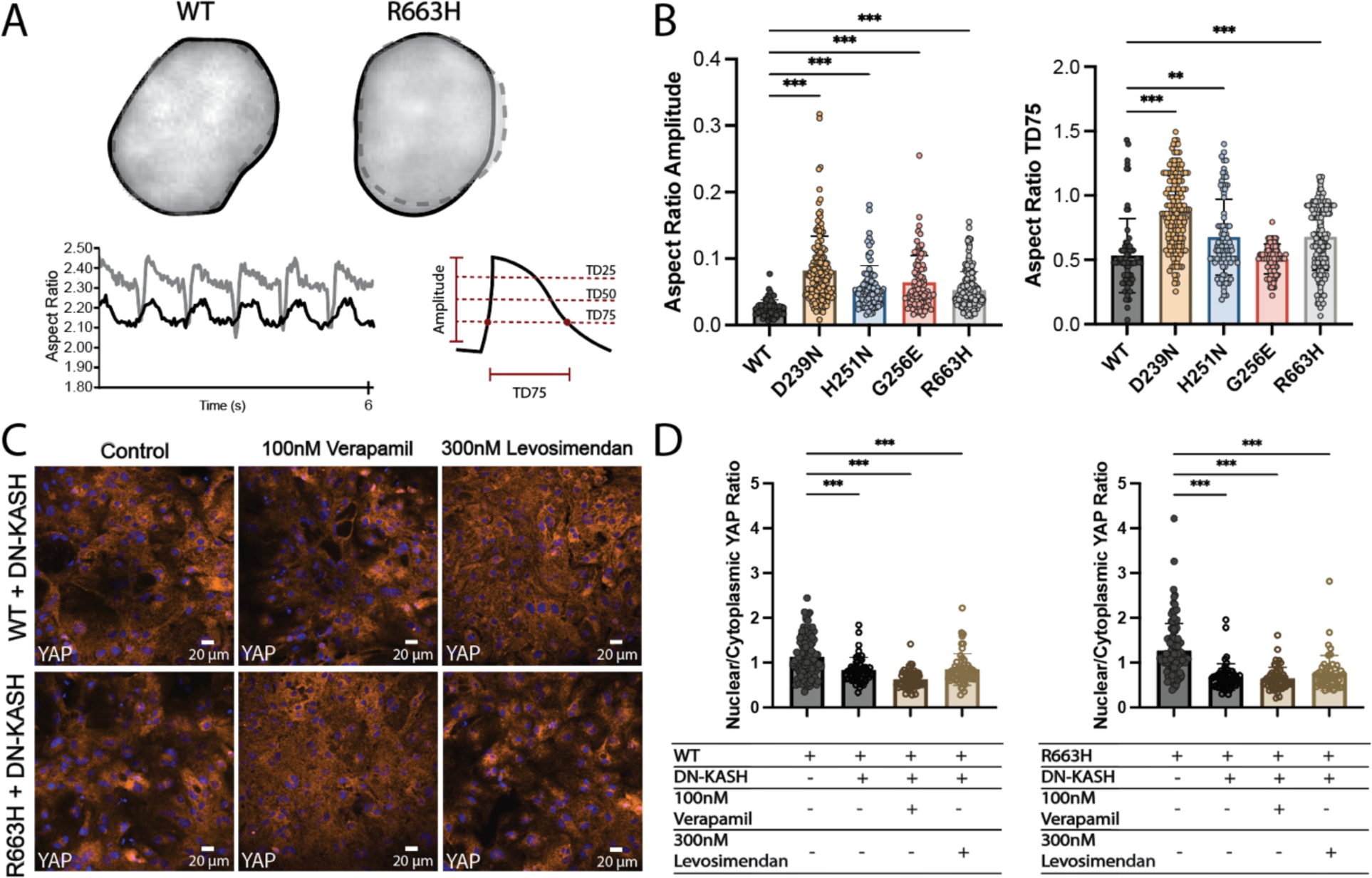
YAP activation in MYH7 mutants requires nuclear mechanical coupling. (A) Outlined representative image of WT and MYH7-R663H hiPSC-CM nucleus. Relaxed state (solid black line) and contracted state (dash grey line). Representative change in aspect ratio trace of paced (1hz) WT(black) and MYH7-R663H (grey) hiPSC-CMs. (B) Quantification of nuclear deformation (aspect ratio amplitude) and time under deformation (aspect ratio TD75). Statistical test: Mann-Whitney. (C) Representative images of fixed and stained D(60) WT and MYH7-R663H hiPSC-CMs after DN-KASH transfection and either verapamil or levosimendan treatment. (D) Quantification of YAP activity in WT (left) and MYH7-R663H (right) from DN-KASH drug experiments inferred by Nuclear to Cytoplasmic intensity Ratio. Statistical test: one-way ANOVA. P<0.05 is designated with (*), P<0.005 is designated with (**). P<0.0005 or smaller is designated with (***).

Next, we inquired whether nuclear deformation and associated increases in YAP nuclear localization could be disrupted by interfering with the LINC complex (thereby inhibiting mechanical strain on the nuclei)^18,24^. Transfecting hiPSC-CMs with a dominant-negative plasmid for nesprin 1 (DN-KASH) impeded the interaction between nesprin and sun proteins to impair cell/nuclear coupling^24^. Both WT and MYH7-R663H mutant hiPSC-CMs were transfected with DN-KASH and successfully incorporated hiPSC-CMs evaluated for YAP localization through immunocytochemistry after a 24-hour incubation period (Figure 4C). In both WT and MYH7-R663H mutant hiPSC-CMs, disruption of the LINC complex led to significantly diminished YAP nuclear localization (Figure 4C-D).

We then tested whether increasing or reducing contractility via positive and negative inotropes in hiPSC-CMs with disrupted LINC complexes, exacerbated or rescued N/C YAP ratio to control levels. Treatment with negative and positive inotropic drugs—verapamil (100 nM) and levosimendan (300 nM)—failed to restore or alter the reduced YAP nuclear localization. To validate alterations in nuclear deformation due to drug treatment and/or LINC complex disruption, nuclear deformation was measured as described previously (Supplement 4D-F). Levosimendan (300 nM) treatment alone on healthy (WT) hiPSC-CMs resulted in enhanced nuclear deformation (aspect ratio amplitude) (Supplement 4E). Conversely, verapamil (100 nM) treatment alone on MYH7-R663H mutant hiPSC-CMs led to reduced nuclear deformation (aspect ratio amplitude) (Supplement 4F). Successful disruption of the LINC complex in hiPSC-CMs resulted in reduced nuclear deformation, even when combined with inotropic drug treatments (Supplement 4E-F). Together, these experiments suggest that (1) contractile force-mediated nuclear deformation influences nuclear entry of YAP (2) disruption of the cytoskeletal/nuclear assembly (LINC complex) can impede YAP nuclear entry by disconnecting the mechanical elements responsible for transmitting mechanical signals to the nuclear membrane.

### Reducing hypercontractility of MYH7-R663H mutant hiPSC-CMs alters the transcriptomic profile to resemble healthy (WT) hiPSC-CMs

We next asked whether transcriptional changes accompany the observed increases in nuclear deformation and YAP nuclear localization. To characterize the transcriptional landscape of MYH7 mutant hiPSC-CMs we assessed global gene expression profiles by conducting bulk RNA sequencing (RNA-seq). We performed bulk RNA-seq on D60 hiPSC-CMs from four separate conditions: *1) WT hiPSC-CMs*, **2) WT hiPSC-CMs + levosimendan (300 nM)** to mimic the hypercontractile phenotype, **3) MYH7-R663H hiPSC-CMs**, and *4) MYH7-R663H hiPSC-CMs+ verapamil (100 nM)* in an effort to reduce hypercontractility and therefore presumably restore a normal transcriptional profile (**Bold** = high force, *Italics* = low force). Initial examination of the transcriptomes of *WT* and **MYH7-R663H** hiPSC-CMs revealed an upregulation of known cardiomyocyte markers ACTN2, and MYH7 (Supplement 5A). **MYH7-R663H** mutant hiPSC-CMs predominantly expressed MYH7 over MYH6, a common association with HCM (Supplement 5A). Additionally, we observed a significant enrichment of hallmark HCM genes NPPA and NPPB, among others, suggesting that our model captures key transcriptomic features expected of HCM (Supplement 5B-C). Gene networks were analyzed through Kyoto Encyclopedia of Genes and Genomes (KEGG) pathway analysis of the top differentially expressed genes (DEGs). Analysis indicated an enrichment in pathways such as “Focal Adhesion”, “ECM Receptor Interaction”, “Hypertrophic Cardiomyopathy”, “Regulation of Actin Cytoskeleton”, “TGFβ Signaling”, and “Hippo Signaling” (Figure 5A)^72^. When all four samples were unbiasedly compared, *WT* and *MYH7-R663H + verapamil (100 nM)* hiPSC-CMs were more similar and **WT + levosimendan (300 nM)** and **MYH7-R663H** hiPSC-CMs were more similar. These similarities are visualized in a hierarchical heatmap and Principal Components Analysis (PCA) plot (Supplement 5D-E). We hypothesize that the variance along the x-axis (PC1) is associated with force generation (low to high), while the variance along the y-axis (PC2) is associated with disease (healthy to disease) (Supplement 5E). We grouped the DEG heatmap into 4 gene clusters (Figure 5B). Gene Cluster 1 appears to be unique to the *WT* line and Gene Cluster 4 unique to the **MYH7-R663H** mutant line. While Gene Cluster 3 includes genes associated with an increase in peak force production by **MYH7-R663H** mutant and **WT + levosimendan** treatment hiPSC-CMs.

**Figure 5.**
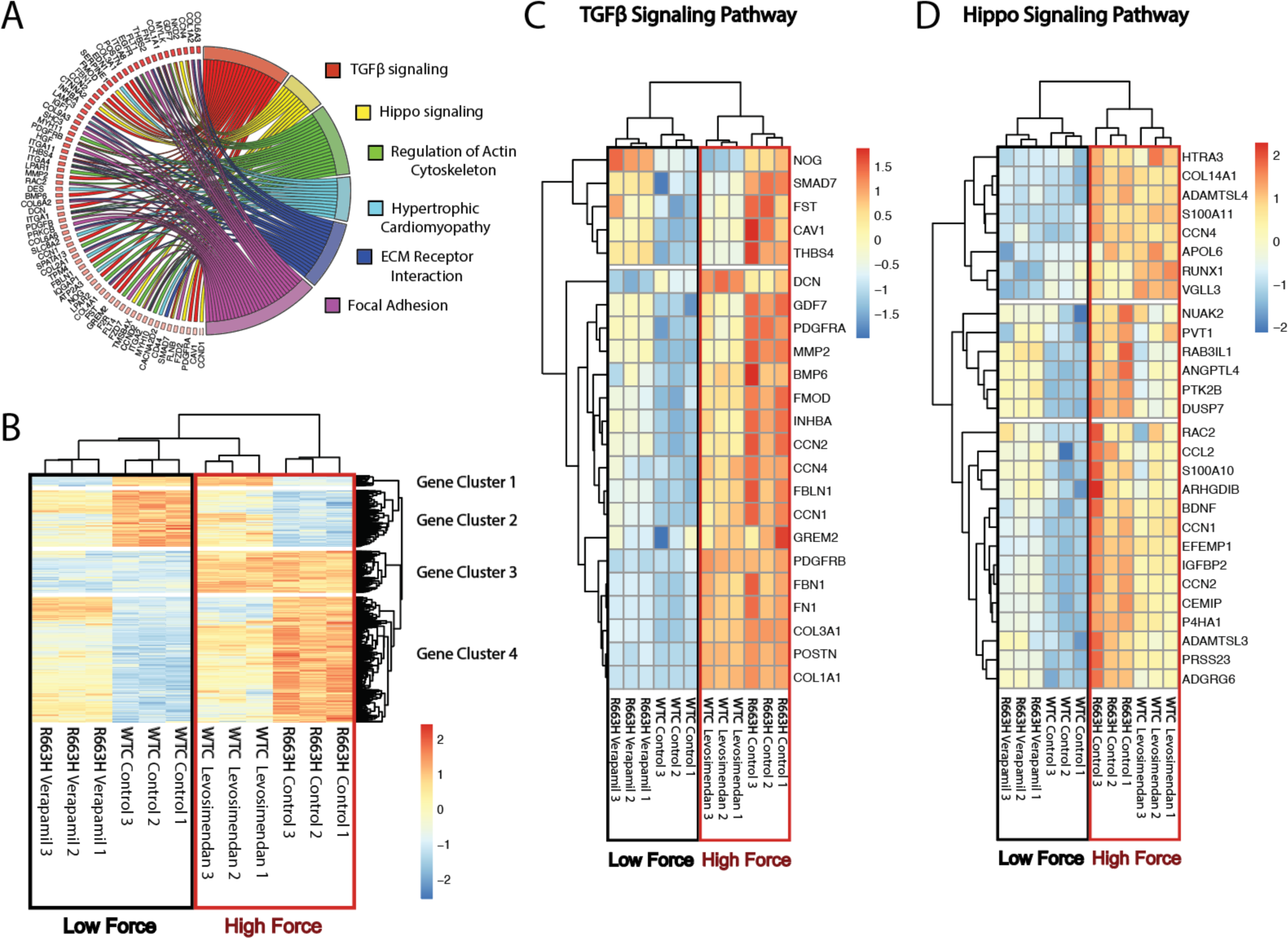
Force modulation dysregulates both Hippo and TGFB1 signaling pathways in WT and MYH7-R663H mutant hiPSC-CMs. (A) Chord plot illustrating the gene ontology of KEGG pathways from the top differentially expressed genes (DEGs) comparing D60 WT and MYH7-R663H mutant hiPSC-CM (fold change > 2, p-value < 0.05). (B) Heat map illustrating the top DEGs hierarchically clustered on similarity and further sub-grouped into four separate identifiable gene clusters. The red box indicates hiPSC-CMs with “High Force” production and black box indicates hiPSC-CMs with “Low Force” production. (C-D) Heat maps depicting force mediated transcriptional change of genes related to TGFβ signaling and Hippo Signaling pathways. The red box indicates hiPSC-CMs with “High Force” production and black box indicates hiPSC-CMs with “Low Force” production.

A subset of these force-related genes appear in KEGG pathways “TGFβ Signaling” and “Hippo Signaling” (Figure 5C-D). Notably, a specific gene family of secreted signaling proteins, Cellular Communication Network (CCN), was upregulated in both “TGFβ Signaling” and “Hippo Signaling” pathways (CCN1, CCN2, and CCN4). CCN2 is a well-known downstream target of YAP and often serves as an indicator of transcriptional YAP activation^28,73^.

### Conditioned media, from either MYH7-R663H mutant hiPSC-CMs or WT hiPSC-CMs with increased contractility, contains increased levels of CCN2 and results in the activation of hiPSC-CFs

CCN2, a paracrine signaling molecule shared between cardiomyocytes and cardiac fibroblasts, has been implicated in cardiac fibroblast/myofibroblast transition and fibrosis initiation^74–77^. Since our RNA-seq suggests upregulation of CCN related protein CCN2, we asked whether cardiomyocytes experiencing hypercontractility activate the CCN2 fibrotic signaling pathway through the nuclear localization of YAP. We examined CCN2 transcript levels in D60 MYH7 mutant hiPSC-CMs and found a more than 5-fold increase in mRNA levels compared to WT hiPSC-CMs (Figure 6A). This observation was further confirmed at the protein level through an Enzyme-Linked Immunosorbent Assay (ELISA) that quantified increased CCN2 protein secretion from MYH7-R663H mutant after 48 hours of culture (Figure 6B).

**Figure 6.**
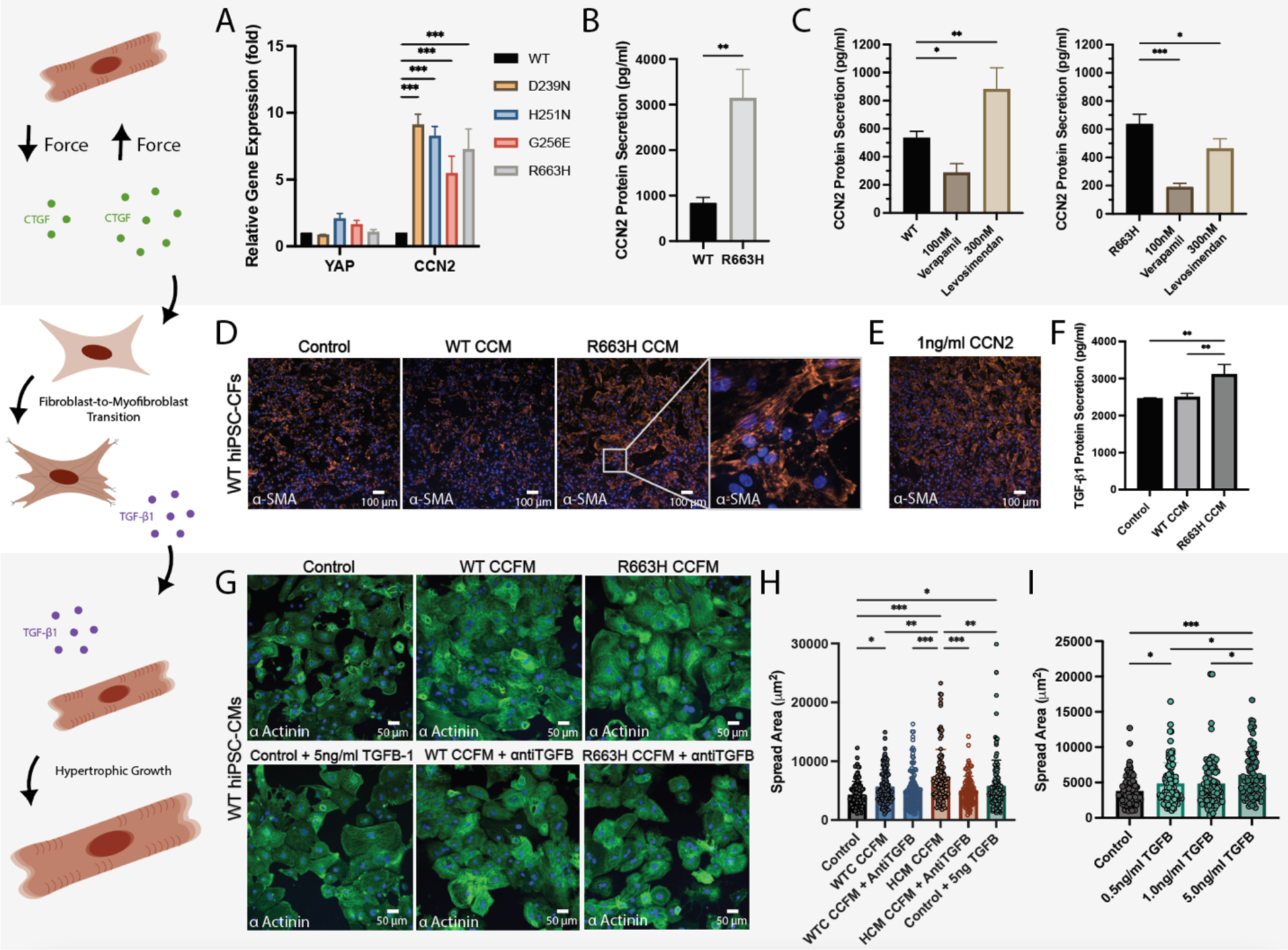
Conditioned media experiments reveal enhanced force production by MYH7-R663H hiPSC-CMs and subsequent increased CCN2 secretion, activates pro-hypertrophic TGFβ secretion by hiPSC-CFs. (A) qPCR results for genes: YAP and CCN2 for all four MYH7 mutants (D239N, H251N,G256E and R663H). Statistical test: Mann-Whitney. (B) Bar plot illustrating CCN2 Elisa results for WT and R663H hiPSC-CMs spent media (48hrs). Statistical test: unpaired t-test between mutant and paired isogenic control. (C) Bar plots depicting CCN2 Elisa results for WT (left) and R663H (right) hiPSC-CMs spent media with 48hr drug treatments (verapamil(100 nM) and levosimendan (300 nM)). Statistic test: one-way ANOVA. (D) representative images of hiPSC-CFs fixed and stained for αSMA (RFP), reveal enhanced activation of hiPSC-CFs after R663H CCM treatment (144hrs). (E) representative images of hiPSC-CFs fixed and stained for αSMA, after 1 ng/ml CCN2 treatments reveals comparable activation of hiPSC-CFs when compared to R663H CCM treatment (144hrs total, 48hr media exchanges). (F) TGFβ Elisa results of hiPSC-CFs after 144hr treatment with WT CCM and R663H CCM. Statistic test: one-way ANOVA. (G) Representative images of WT hiPSC-CMs fixed and stained for αactinin (GFP), after 144hr treatment with double conditioned media WT CCFM and R663H CCFM, TGFβ blocking antibody (anti TGFβ), and purified human TGFβ (5 ng/ml). (H) quantification of WT hiPSC-CM spread area after the above mentioned treatments. Statistic test: Kruskal-Wallis. (I) quantification of WT hiPSC-CM spread area after purified human TGFβ −1 treatment. Statistical test: Kruskal-Wallis. P<0.05 is designated with (*), P<0.005 is designated with (**). P<0.0005 or smaller is designated with (***).

We next wanted to corroborate the connection between force, YAP nuclear localization, and CCN2 transcriptional activation supported by bulk RNA-seq results. We performed an ELISA for CCN2 protein on WT and MYH7-R663H hiPSC-CMs with and without negative inotrope verapamil (100 nM) and positive inotrope levosimendan (300 nM) treatment for 24 hours (Figure 6C). The results demonstrated significantly depleted CCN2 production in force reduced verapamil-treated WT hiPSC-CMs and increased CCN2 production in force enhanced levosimendan-treated WT hiPSC-CMs. Suggesting a regulatory role of cardiomyocyte contraction on CCN2 secretion. Treatment of MYH7-R663H mutant hiPSC-CMs with verapamil (100 nM) significantly decreased CCN2 production. However, levosimendan treatment led to significantly reduced CCN2 production in MYH7-R663H hiPSC-CMs, consistent with its confounding effects observed on YAP nuclear localization in Figure 3B.

We assessed whether driving YAP localization has a direct influence on CCN2 secretion by utilizing TRULI, a small molecule shown to upregulate YAP localization and downstream targets^78–80^. When WT hiPSC-CMs were treated with TRULI (2 µM) for 24 hrs we observed significantly increased nuclear localization of YAP, CCN2 mRNA expression and CCN2 protein secretion, thus supporting the role of YAP and CCN2 secretion in HCM mutant hiPSC-CMs (Supplement 6A-B). However, when we attempted to alternatively inhibit YAP activation neither Verteporfin, a well-documented YAP inhibitor demonstrated in other cell types, or TM2, (R)-PFI-2, and TAT_PDHPS1 resulted in decreased YAP activation nor CCN2 secretion (Supplement 6C-F)^81–84^.

Previous studies have demonstrated the propensity of secreted CCN2 to activate cardiac fibroblasts and trigger the fibroblast to myofibroblast transition^6,75,85,86^. We further investigated whether the abundance of CCN2 in spent media from MYH7-R663H mutant hiPSC-CMs could activate hiPSC-derived cardiac fibroblasts (hiPSC-CFs), inducing their transition to cardiac myofibroblasts defined by elevated levels and incorporation of alpha-smooth muscle actin (αSMA) into stress fibers^87,88^. Spent media (48-hour culture) from either D60 WT or MYH7-R663H mutant hiPSC-CMs were mixed 1:1 with fresh cardiac fibroblast media (referred to as conditioned cardiac media or CCM, illustration of experimental design in Supplement 7). WT hiPSC-CFs were subjected to treatment with both WT CCM and MYH7-R663H CCM (144 hours in total, with media exchanged every 48 hours), followed by fixation and staining for alpha-smooth muscle actin (αSMA). We observed an increase in hiPSC-CFs transitioned to cardiac myofibroblasts, evidenced by an increase in αSMA intensity and incorporation into stress fibers (Figure 6D, Supplement 8A). To verify that this enrichment of αSMA was specifically triggered by secreted CCN2, WT hiPSC-CFs were treated with purified human CCN2 at a concentration of 1 ng/ml. We observed a similar enhancement in αSMA intensity and incorporation into stress fibers (Figure 6E, Supplement 8A). Collectively, these findings suggest that CCN2-enriched MYH7-R663H mutant media activates the transition of cardiac fibroblasts to myofibroblasts, implicating its potential role in fibrotic signaling.

### Cardiac fibroblasts treated with CCN2 enriched MYH7-R663H mutant media, subsequently secrete a known hypertrophic signaling molecules TGFβ

Next, we explored whether cardiac myofibroblasts treated with CCN2-enriched MYH7-R663H mutant media secrete known hypertrophic signaling molecule Transforming Growth Factor-β (TGFβ), as described in previous studies^32,34^. Among the three isoforms of TGFβ (TGFβ1, TGFβ2, TGFβ3), TGFβ1 has been widely implicated in various tissue types and has been observed to be upregulated during cardiac hypertrophy and remodeling^89,90^. We investigated whether activated cardiac myofibroblasts, described in the previous experiment, subsequently induce the secretion of TGFβ1^91^. Following 144 hours of treatment with WT-CCM or MYH7-R663H-CCM (media exchanged every 48 hours), we collected spent media from hiPSC-CFs, and performed a TGFβ1 ELISA. We saw a significant increase in the amount of secreted TGFβ1 in the spent media of hiPSC-CFs treated with MYH7-R663H CCM compared to WT CMM treated and control hiPSC-CFs (Figure 6F). Previous research has highlighted TGFβ1 as a homeostatic signaling molecule responsible for cardiac tissue injury repair and remodeling^90^. Specifically, TGFβ1 has been shown to further induce the transition of cardiac fibroblasts to myofibroblasts through the reactivation of CCN2 production and acts as a feedforward pro-hypertrophic signal on cardiomyocytes (Supplement 8B)^89,92^. Finally, we investigated whether the enriched TGFβ1 double-conditioned media (CCM + hiPSC-CF spent media) affected cardiomyocyte hypertrophy, as assessed by changes in spread area^39^. We pivoted from the use of a complex co-culture experiment to a reduced order conditioned media experiment due to (1) the sensitivity of hiPSC-CM to stressors (i.e. glucose starvation) and the subsequent confounding effects on cell size, and (2) the overgrowth of cardiac fibroblasts/myofibroblasts in extended cultures inhibiting cardiomyocyte function (illustration of experimental design in Supplement 7)^93,94^. Since TGFβ1 has been shown to play a significant role in mediating the hypertrophy of cardiomyocytes, we evaluated whether the conditioned medium from CCM-treated hiPSC-CFs (referred to as conditioned cardiac fibroblast media (CCFM)) affected hiPSC-CM cell size (Supplement 7). We measured the spread area of hiPSC-CMs after 144 hours of CCFM treatment. We observed an increase in the spread area of WT hiPSC-CMs when treated with doubly conditioned hiPSC-CM WT CCFM media compared to control, with an even greater increase observed when WT hiPSC-CMs were treated with doubly conditioned hiPSC-CM MYH7-R663H CCFM media (Figure 6G-H). To further verify if TGFβ1 was directly responsible for the enlargement of hiPSC-CMs, we utilized a TGFβ inhibiting antibody (antiTGFβ, blocking TGFβ1, TGFβ2, and TGFβ3) to block signaling and purified human TGFβ1 to directly activate signaling. Anti-TGFβ inhibiting antibody was supplemented to the doubly conditioned CCFM treatments at the manufacturer’s recommended dose of 3.75 ng/ml (Figure 6H, Supplement 8C). Interestingly, while blocking TGFβ1 did not affect the spread area of WT-CCFM-treated WT hiPSC-CMs, it did significantly reduced the spread area change in hiPSC-CMs treated with MYH7-R663H CCFM. Moreover, the addition of 5 ng/ml of purified human TGFβ1 to WT hiPSC-CMs expectedly increased the spread area (Figure 6H-I). However, the size of purified (glucose-starved) WT and MYH7 mutant hiPSC-CMs remained unchanged (Supplement 8D). These observations confirm the importance of hiPSC-CFs and potentially other cell types in driving the manifestation of disease phenotypes. Collectively, these data suggest a paracrine signaling interaction between hiPSC-CMs and hiPSC-CFs may contribute to the hypertrophic growth and fibrotic signaling observed in HCM.

## Discussion

While inherited cardiovascular diseases continue to pose a significant health challenge, our understanding of how small genetic alterations can result in larger life threatening complications remains incomplete. Several significant limitations have impeded our comprehension of cardiovascular diseases like HCM, including: (1) the scarcity of human primary cardiomyocytes, (2) the inability to culture human primary cardiomyocytes for prolonged periods, and (3) discrepancies between animal and human models, especially in sarcomeric protein isoforms. In this study we employ the power of hiPSC-derived cardiomyocytes as a human model to explore the role of YAP activation in HCM. While previous groups have described the activation of YAP in human HCM tissue or hypertrophic mouse models (TAC), the extent of their investigation on YAP’s involvement in the manifestation of HCM remains modest^31^.

We confirm elevated YAP activation in hiPSC-CMs and human tissue harboring HCM-associated MYH7 mutations (Figure 1A-C). This marks the first verification that YAP activation in human cardiac tissue is selective to the cardiomyocyte population, a detail missed by previous studies due to their chosen methodologies when investigating diseased tissue (Western Blot or RNA of bulk tissue samples, Figure 1E)^30,31^. Additionally, we observe no alterations in YAP isoform expression in HCM mutant hiPSC-CMs (Supplement 1A). With the confirmation of altered YAP activity in HCM-associated MYH7 mutant hiPSC-CMs and human cardiac tissue, we set out to answer our three original questions:

### How does enhanced contractility alter the transcriptomic behavior of cardiomyocytes?

We suspected that the preliminary phenotype of hypercontractility drives the subsequent hypertrophic and fibrotic phenotypes of HCM. We confirm that hypercontractility is accompanied by increased YAP activity (nuclear localization) in four MYH7 mutant hiPSC-CMs (Figure 2A-C). To test whether hypercontractility alone is sufficient to increase YAP activity, we use positive and negative inotropic drugs (levosimendan and verapamil, respectively), and increased substrate stiffness (Supplement 2D-G). Our detailed studies focus on the MYH7-R663H mutant because of the availability of a confirmatory patient sample. The results reveal that increasing force generation in WT hiPSC-CMs is accompanied by the nuclear accumulation of YAP, and decreasing force generation in MYH7-R663H and WT hiPSC-CMs depletes levels of nuclear YAP, suggesting contractility as a mediator of YAP’s nuclear localization (Figure 3A-B). We characterize the transcriptome of healthy (WT) and diseased (MYH7-R663H) hiPSC-CMs and demonstrate the upregulation of well-known HCM genes MYH7, NPPA, and NPPB, demonstrating the fidelity of our model (Supplement 5A-C). Treatment with inotropic drugs to either induce a hypercontractile state of a WT hiPSC-CM (WT + levosimendan) or rescue the hypercontractile phenotype of the MYH7-R663H mutant hiPSC-CM (MYH7-R663H + verapamil) suggested a host of genes correlate with contractile force (Figure 5B). KEGG pathways analyses of WT, WT + levosimendan, MYH7-R663H, and MYH7-R663H + verapamil suggested the upregulation of genes closely related to both the Hippo and TGFβ pathways (Figure 5C-D). Further exploration revealed that part of the CCN protein family was upregulated. The CCN protein family of genes, specifically CCN2, is modulated by YAP^28,73^ and initiates fibrotic signaling during cardiac injury^74–77^.

### How does a cardiomyocyte modulate YAP activity in a mechanically dynamic environment?

Utilizing a uniaxial stretch device we demonstrated the mechanics of contractility to be the predominant driver of YAP nuclear localization (Supplement 3A-C). Although the mechanical activation of YAP has been well studied, more recent studies suggest YAPs nuclear localization is driven by deformations of the nucleus^24^. Our findings complement and confirm a correlation between YAP nuclear localization, hypercontractility and nuclear deformation for all four HCM mutant hiPSC-CM lines (Figure 4A-C). By inhibiting the mechanical deformations of the nucleus through the inhibition of the LINC complex (DN-KASH), we were able demonstrate the dependence of YAP nuclear localization on the deformation of the nucleus (Figure 4C-D). The upregulation of nuclear structural proteins suggests a compensation mechanism used by hiPSC-CMs to compensate for the increase in nuclear deformation driven by increased contractile force (Supplement 4C). We hypothesized the hypercontractility phenotype of HCM drives the activation of YAP through the physical deformation on the nucleus.

### Lastly, To what extent do these alterations contribute to the phenotypic manifestations of hypertrophic cardiomyopathy?

After confirming elevated levels of CCN2 RNA and protein, we tested the effects of CCN2 on hiPSC-CFs (Figure 6A-B). Subjecting hiPSC-CFs to spent HCM mutant hiPSC-CM media induced the activation of cardiac myofibroblast, demonstrated by the enhanced levels and incorporation of αSMA into stress fibers (Figure 6D). As a result of myofibroblast activation, further analysis depicted enhanced levels of known secreted hypertrophic signaling molecule TGFβ by hiPSC-CFs (Figure 6F).

Altogether, our results strongly support a model in which paracrine signaling exists between hypercontractile cardiomyocytes and cardiac fibroblasts. Specifically, CCN2 mediated myofibroblast activation and subsequent increased cardiac fibroblast TGFβ secretion leads to the hypertrophic growth and fibrotic remodeling of the heart in HCM.

## Study Limitations

Our study utilizes hiPSC-derived cell types, which are known to be immature and do not fully recapitulate all features of human heart tissue in vivo. However they have been shown to be suitable to model disease processes especially at the cellular level^39–44^. The single-cell, reduced-ordered model described here has provided measurements that otherwise would not be possible. Other limitations include, but are not limited to, differences introduced between sexes, multiple parental cell lines, or patient tissue with the same mutations.

## Summary

Our research has demonstrated the following: (1) Confirmed that MYH7 mutations D239N, H251N, G256E lead to hypercontractility in hiPSC-derived cardiomyocytes (hiPSC-CMs); (2) Increase in YAP’s nuclear localization in both hiPSC-CMs and patient tissue with MYH7 HCM mutations; (3) A correlation between YAP’s nuclear localization and enhanced force generation; (4) Nuclear deformation as a mechanism enabling YAP’s nuclear entry and activation; (5) Transcriptomic changes resulting from positive and negative modifications in force generation; and (6) A distinctive paracrine hypertrophic signal reliant on cardiomyocyte-cardiac fibroblast crosstalk. Our unique insights to the intricacies of hypertrophic cardiomyopathy phenotypes hold the potential to provide mechanistic clarity, informing potential therapeutic strategies.

## Methods

### Maintenance of stem cell culture

The human induced pluripotent stem cell (hiPSC) parental line WTC, was generated by the Bruce R. Conklin Laboratory at the Gladstone Institutes and University of California-San Francisco (UCSF)^95^. hiPSC lines MYH7-G256E, MYH7-H251N, and MYH7 isogenic controls WT_G256E_ and WT_H251N_ were developed at the Allen Institute and shared with the Beth Pruitt lab at the University of California Santa Barbara (UCSB) as part of greater collaboration with the University of Washington, Stanford University and UCSB. These lines were created in an established αActinin-GFP WTC background. The MYH7-D239N hiPSC line and its isogenic control WT_D239N_ were gifted from the Daniel Bernstein lab at Stanford University. The MYH7-R663H patient derived hiPSC line was obtained from the Stanford Cardiovascular Institute (SCVI) Biobank^96^. To serve as a control for the unique patient-derived cell line (MYH7^WT/R663H^), the parental unedited WTC (GM25256) hiPSC line was used. Cell line details and reference names can be found in Table 1.

hiPSCs were propagated on Matrigel® coated plates (dilution 1:200) using feeder-free culture conditions (mTeSR™ and Essential 8) in standard environments consisting of 5% carbon dioxide at 37°C. Media was replaced daily and cells were passaged with EDTA when hiPSC cultures reached 80% confluence.

| Parental Cell Line | HCM Mutation | Reference Name |
| --- | --- | --- |
| SCVI-113 | MYH7<sup>WT/D239N</sup> | D239N |
| SCVI-113 | MYH7<sup>WT/WT</sup> | WT<sub>D239N</sub> |
| WTC (AICS-75 cl85) | MYH7<sup>WT/H251N</sup> | H251N |
| WTC (AICS-75 cl85) | MYH7<sup>WT/WT</sup> | WT<sub>H251N</sub> |
| WTC (AICS-75 cl85) | MYH7<sup>WT/G256E</sup> | G256E |
| WTC (AICS-75 cl85) | MYH7<sup>WT/WT</sup> | WT<sub>G256E</sub> |
| Patient Line | MYH7<sup>WT/R663H</sup> | R663H |
| WTC(GM25256) | MYH7<sup>WT/WT</sup> | WT<sub>R663H</sub> |

### hiPSC-CM and hiPSC-CF differentiation

Directed cardiac differentiation was achieved using the WNT modulating protocol previously described^51^. Upon D12 of cardiac differentiation, hiPSC-CMs were purified by glucose starvation (RPMI(-Glucose) + B27 supplement) as previously described^97^. hiPSC-CMs were further maintained in RPMI + B27 supplement with media changes every 48hrs.

Directed cardiac fibroblast differentiation was achieved using the previously described protocol^98^. Upon D32 of cardiac fibroblast differentiation, hiPSC-CFs were passaged and maintained in Promocell Fibroblast Growth Medium 3 with media changes every 48 hrs. hiPSC-CFs were passaged with Accutase® when cultures reached 80% confluence.

### Traction Force Microscopy

Traction Force Microscopy was conducted following established procedures^99^. Single micropatterned hiPSC-CMs were imaged using a Zeiss Axio Observer 7 inverted microscope equipped with a high speed/resolution camera (Photometrics Prime 95b). Micropatterns of 1500µm^2^ area with a 7:1 aspect ratio were created using microcontact printing. Patterned hiPSC-CMs were cultured on polyacrylamide hydrogels with of different stiffness (10kPa, 35kPa or 100kPa) with embedded fluorescent beads (FluoSphere™ Carboxylate-Modified Microspheres, 0.5µm, red fluorescent 580/605) for 48hrs. Displacements of the imaged fluorescent beads over time (i.e. velocity) were quantified using custom Matlab code. Contractile force dipoles were integrated from traction stresses as previously described^99^.

### hiPSC-CM/hiPSC-CFs immunocytochemistry and Human Tissue immunohistochemistry

hiPSC-CMs and hiPSC-CFs were fixed with 4% paraformaldehyde (PFA). Primary antibodies (1:200) and secondary antibodies (1:500) were diluted in PBS supplemented with 0.1% Tween20 (PBST) and 1% BSA. Primary antibodies directed against αActinin (Sigma-Aldrich A7811), YAP (Abcam-52771), and α-SMA(Abcam-124964) were incubated overnight at 4°C. Secondary Alexa 488 or 564 conjugated antibodies were incubated for 30mins and counterstained with Hoechst and CellMask™ (Invitrogen, deep red) prior to imaging.

WT and MYH7-R663H human cardiac tissue was gifted by the Daniel Bernstein lab. Flash frozen human tissue samples were immediately immersed in OCT freezing compound (Fisher), frozen and sectioned using a microtome. Tissue sections were placed on glass microscope slides and stored in the −80°C. Prior to immunohistochemistry, sections were removed from the −80°C and immediately fixed with 4% paraformaldehyde (PFA). The fixed tissue sections were permeabilized and blocked for 1hr in PBS supplemented with 0.3% Tween20 and 2% BSA prior to overnight primary antibody incubation. Tissue sections were then washed and incubated with 0.1% (diluted in ETOH) Sudan Black for 10mins (to mitigate autofluorescence), washed and incubated with secondary antibody for 30mins. Finally, fixed and stained tissue sections were mounted using ProLong™ Gold Antifade with DAPI (Thermofisher) prior to imaging. Negative controls for immunohistochemistry included the use of primary and secondary antibodies alone.

### Imaging experiments

All fixed cell imaging experiments were carried out on a Nikon W2 SoRa spinning disk confocal microscope. Glass-bottom or Polymer-like glass bottom dishes were pretreated with Matrigel (1:200 dilution) and D55 hiPSC-CMs were allowed to settle at least 5 days prior to fixation and staining (D60).

### RNA extraction and quantitative RT-PCR (qPCR)

RNA extraction and purification was performed using the RNeasy mini kit (Qiagen) according to manufacturer’s instructions. Reverse transcription was performed with a total of 300ng of RNA using High Capacity RNA-to-cDNA kit (Invitrogen) according to manufacturer’s instructions. Quantitative RT-PCR (qPCR) was performed using PowerUp™ SYBR™ Green Master Mix (Applied biosystems). Three biological replicates and two technical replicates per sample per gene were performed. Technical replicate cycle threshold (Ct) values were averaged (Average Ct) then normalized to glycerol phosphate dehydrogenase (GAPDH) (ΔCt). ΔCt values were further normalized to the control (WT) (ΔΔCt). Final values are displayed as fold changes (2^(-ΔΔCt)^).

### Westerns and ELISA

hiPSC-CMs were first lysed in RIPA lysis buffer (ThermoFisher) supplemented with Halt™ Phosphatase (78420) and Protease (87786) inhibitor cocktails. Lysates were spun down at 10,000 RPM and supernatant collected for protein quantification using a Pierce™ BCA Protein Assay Kit (ThermoFisher-23225). NuPAGE™ 4% to 12%, Bis-Tris mini protein gels (NP0321) were loaded with 15µg of protein for each sample and run in NuPAGE™ MOPS SDS running buffer (ThermoFisher). Gels were then transferred to a PVDF membrane using Pierce™ 1-step transfer buffer (84731). Primary antibodies (1:1000) directed against YAP (Cell Signaling-4912), pYAP S127 (Cell Signaling-4911), LATS1/2 (Bioss-BS-4081R), pLATS1 T1079+ pLATS2 T1041 (Abcam-111344), MST1 (Cell Signaling-3682), pMST1/2 (Cell Signaling 49332) and GAPDH (Proteintech-60004-1) were incubated overnight at 4°C. Secondary LI-COR 680 or 800 conjugated antibodies were incubated for 1hr and imaged on a LI-COR 9120 Odyssey Infrared imaging system. Human CCN2 (Abcam-261851) and TGFβ1 (Abcam-100647) ELISAs were run according to manufacturer’s instructions. ELISA plate preparations were then read on a Synergy H1 microplate reader (BioTek).

### Study cohort and myocardial sample collection HCM patients

The HCM patient underwent septal myectomy for clinical indications at Stanford Medical Center. Inclusion criteria: normal or hyperdynamic LV function (left ventricular ejection fraction (LVEF) > 55%) with either eccentric or concentric LV hypertrophy on echo and a gradient across the left ventricular outflow tract (LVOT). The patient did not have prior evidence of myocardial infarction, primary valvular disease or had progressed to LV dysfunction. A family history of cardiac related disease was obtained and defined as the presence of one or more affected family members with either HCM, arrhythmia, sudden cardiac death, or heart failure. The Stanford Institutional Review Board approved the study.

Controls: Myocardial tissue from both interventricular septum and left ventricular apex were obtained from a donor heart with no major cardiac history, maximum travel distance to Stanford <60 miles, donor ischemic time <6 hours and no history of blunt chest trauma. Cardiac tissue was excised, and a mid-myocardial portion was used immediately for studies of mitochondrial respiration or fixed in 4% paraformaldehyde (PFA) for paraffin embedding or in 4% PFA and 2% glutaraldehyde for TEM analysis. The remaining tissue was flash frozen in liquid nitrogen for all other assays.

### Histological assessments

Fresh myocardial tissue samples (HCM; control) were washed with normal saline solution followed by fixation in 4% paraformaldehyde (PFA) for 24h. The tissue was then embedded in paraffin and sectioned to 7um thickness. After deparaffinization, adjacent sections were stained with hematoxylin and eosin (H&E) to assess tissue morphology and Masson trichrome for collagen deposition associated with fibrosis. This histology was performed by Histotec. All images were taken with a Keyence microscope (BZ-9000, Keyence, Osaka, Japan).

### Small Molecules and inhibiting antibody

Small Molecules: verapamil (Tocris-0654), levosimendan (Sigma-Aldrich-L5545), blebbistatin (Sigma-Aldrich-B0560) and inhibiting antibody: TGFβ-1,2,3 (R&Dsystems-MAB1835) were resuspended according to manufactures protocol. Appropriate dilutions were made fresh in RPMI supplemented with B27. Control comparison groups were treated with either matching volumes of DMSO or water when appropriate.

### Nuclear Deformation

hiPSC-CMs were treated with Hoechst for 1hr, paced at 1hz, and imaged on a Zeiss Axio Observer 7 inverted microscope with a high speed/resolution camera (Photometrics Prime 95b). The captured videos were processed with an custom ImageJ macro script where Hoechst positive nuclei were selected for aspect ratio measurement over time (20s). The aspect ratio traces were averaged and analyzed with a Matlab script to determine their amplitude, rise time, decay time and TD75.

### RNA sequencing

Samples designated for transcriptome analysis were harvested from D60 hiPSC-CMs. hiPSC-CM RNA was harvested and total RNA was purified as described above. Total RNA was sent to Novogene (Beijing, China) for library construction and 150 base pair pair-end sequencing. The resulting raw FASTQ files were trimmed using the default settings of TrimGalore in paired end mode. Afterwards, STAR (STAR aligner version 2.7.8a) was used to map and quantify gene expression levels using the human genome assembly GRCh37 (hg19). DESeq2 with adaptive shrinkage was used for normalization and differential gene expression calling^100,101^. A shrunken absolute log fold change cutoff of 1 and adjusted p-value cutoff of 0.05 was used for differential gene expression calling. For visualization and clustering, variance stabilizing transformation was performed. The removeBatchEffect function in the LIMMA package was used to remove batch effects observed between two successive rounds of sequencing^102^. David Bioinformatics Resources was used for gene ontology analysis^103^.

### Statistics

Data are presented as mean ± standard deviation (SD). Normality was determined by the Anderson-Darling test or Shapiro-Wilk test and the appropriate parametric or non-parametric statistical tests were performed (listed in figure caption). P<0.05 is designated with (*), P<0.005 is designated with (**). P<0.0005 or smaller is designated with (***).

## Author Contributions

Conceptualization, O.C. and B.L.P.; Methodology, O.C., M.A.F., and M.M.; Investigation, O.C., M.A.F., M.M., Z.S., C.M., J.P., A.C., A.V.R., A.G., T.P., K.V.L., B.R., and J.E.S.; Data Curation, O.C., M.A.F., M.M., Z.S., C.M., A.C., A.V.R., T.P.; Writing-Original Draft, O.C.; Writing-Review & Editing, O.C. M.A.F., M.M., D.B. and B.L.P.; Visualization, O.C. and A.C.; Supervision, R.N.G., N.J.S., D.L.M., D.B., S.S.D., S.J.S., D.O.C, M.Z.W. and B.L.P.; Funding Acquisition, O.C., N.J.S, A.G., D.B. and B.L.P

---

**Supplement 1.**
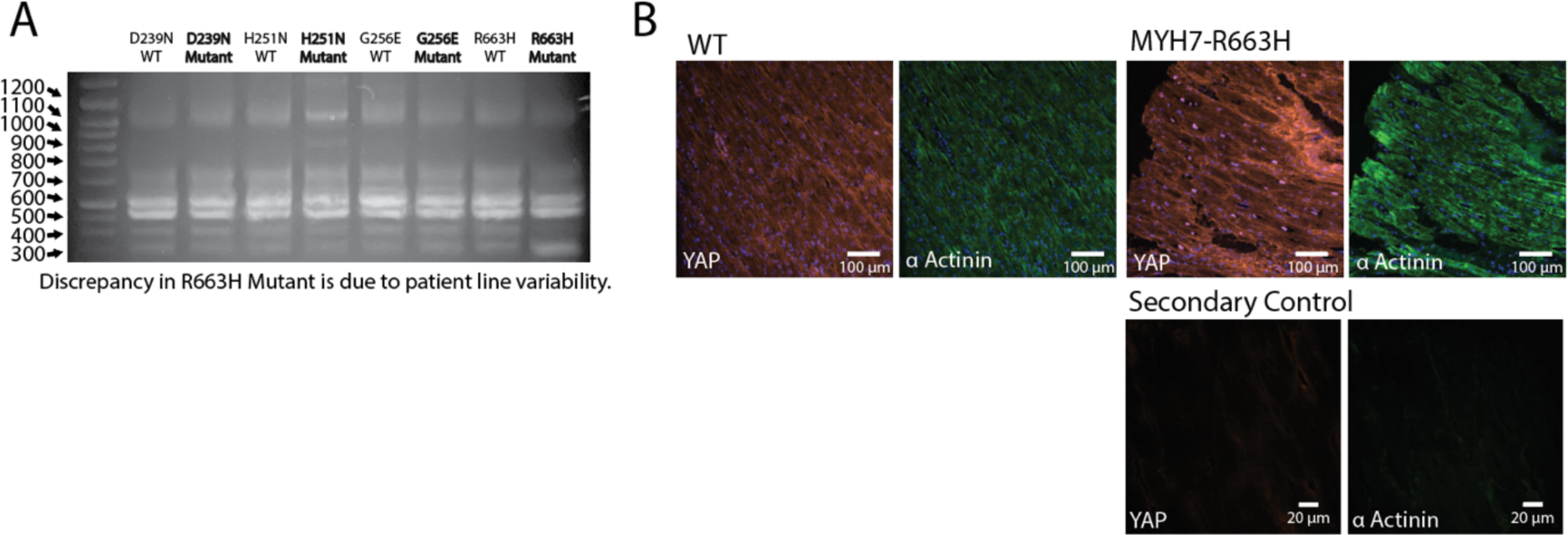
Similar YAP isoform expression in isogenic WT hiPSC-CM and HCM associated MYH7 mutant hiPSC-CMs. (A) PCR amplification of cDNA from each MYH7 mutant hiPSC-CMs and its paired isogenic control. Primer pairs that recognizes conserved portions of the n-and c terminus were used to YAP specific isoforms. (B) Representative image of fixed and sectioned WT and mutant MYH7-R663H human heart tissue labeled for YAP (red), alpha actinin (green) and nuclei (blue) evaluated by Immunohistochemistry. Representative image of fixed and sectioned mutant MYH7-R663H human heart tissue treated with secondary antibody Alexa Fluor 546 (YAP) and Alexa Fluor 488 (α Actinin) only (secondary control), evaluated by Immunohistochemistry.

**Supplement 2.**
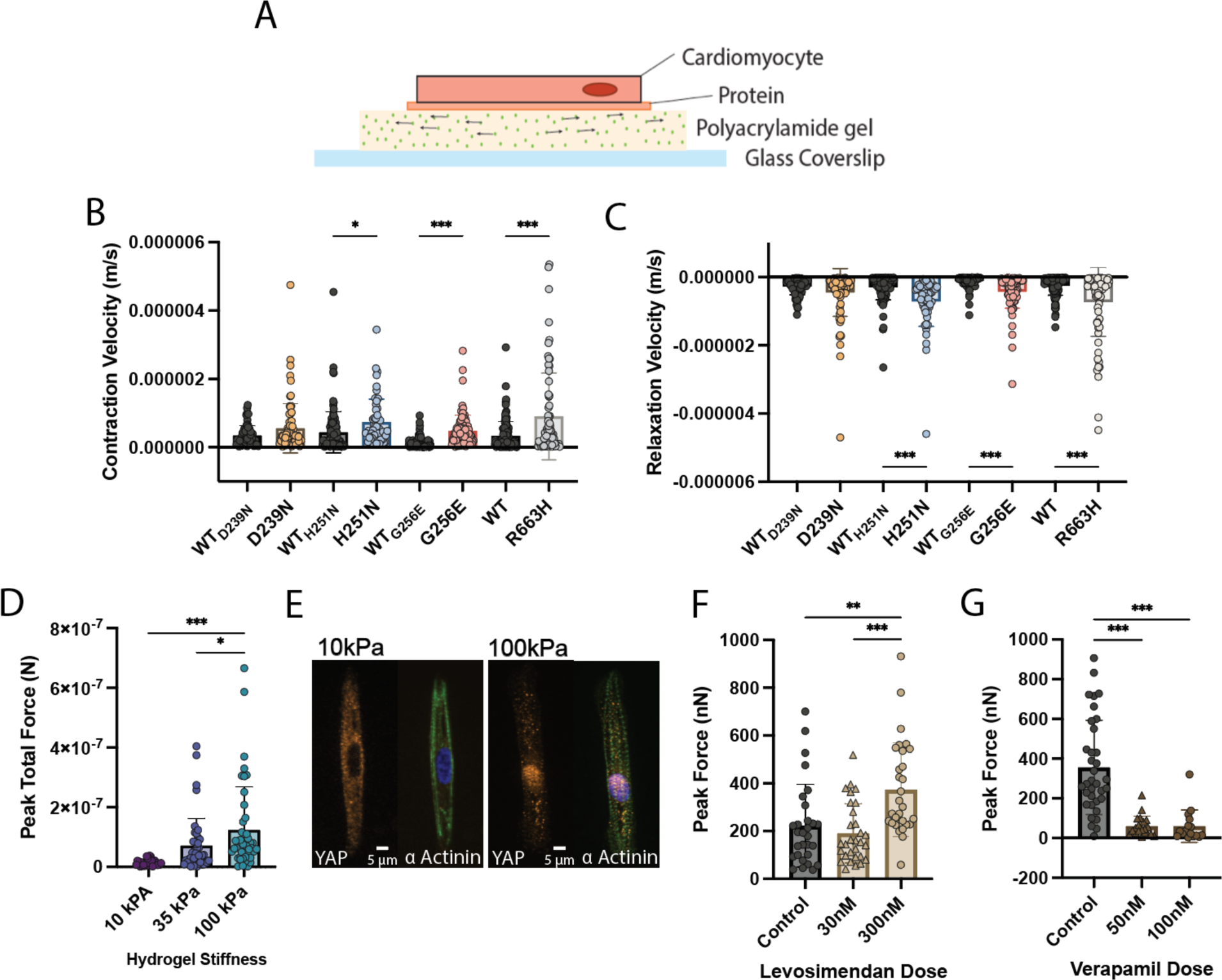
MYH7 mutant hiPSC-CMs develop hypercontractile features. (A) Illustration of the TFM Platform used to assess single patterned hiPSC-CM contractility. (B) Quantification of contraction velocity for each MYH7 mutation and its paired isogenic control. Statistical test: Kruskal-Wallis. (C) Quantification of relaxation velocity for each MYH7 mutation and its paired isogenic control. Statistical test: Kruskal-Wallis. (D) TFM results for WT hiPSC-CMs cultured for 48hrs on 10kPa, 35kPa, and 100kPa polyacrylamide gels. Statistical test: Kruskal-Wallis. (E) Representative images of fixed and stained single patterned hiPSC-CMs cultured for 48hrs on 10kPa and 100kPa polyacrylamide gels. (F) TFM results for WT hiPSC-CMs treated for 24hrs with different levosimendan doses. Statistical test: Kruskal-Wallis. (G) TFM results for WT hiPSC-CMs treated for 24hrs with different verapamil doses. Statistical test: Kruskal-Wallis.

**Supplement 3.**
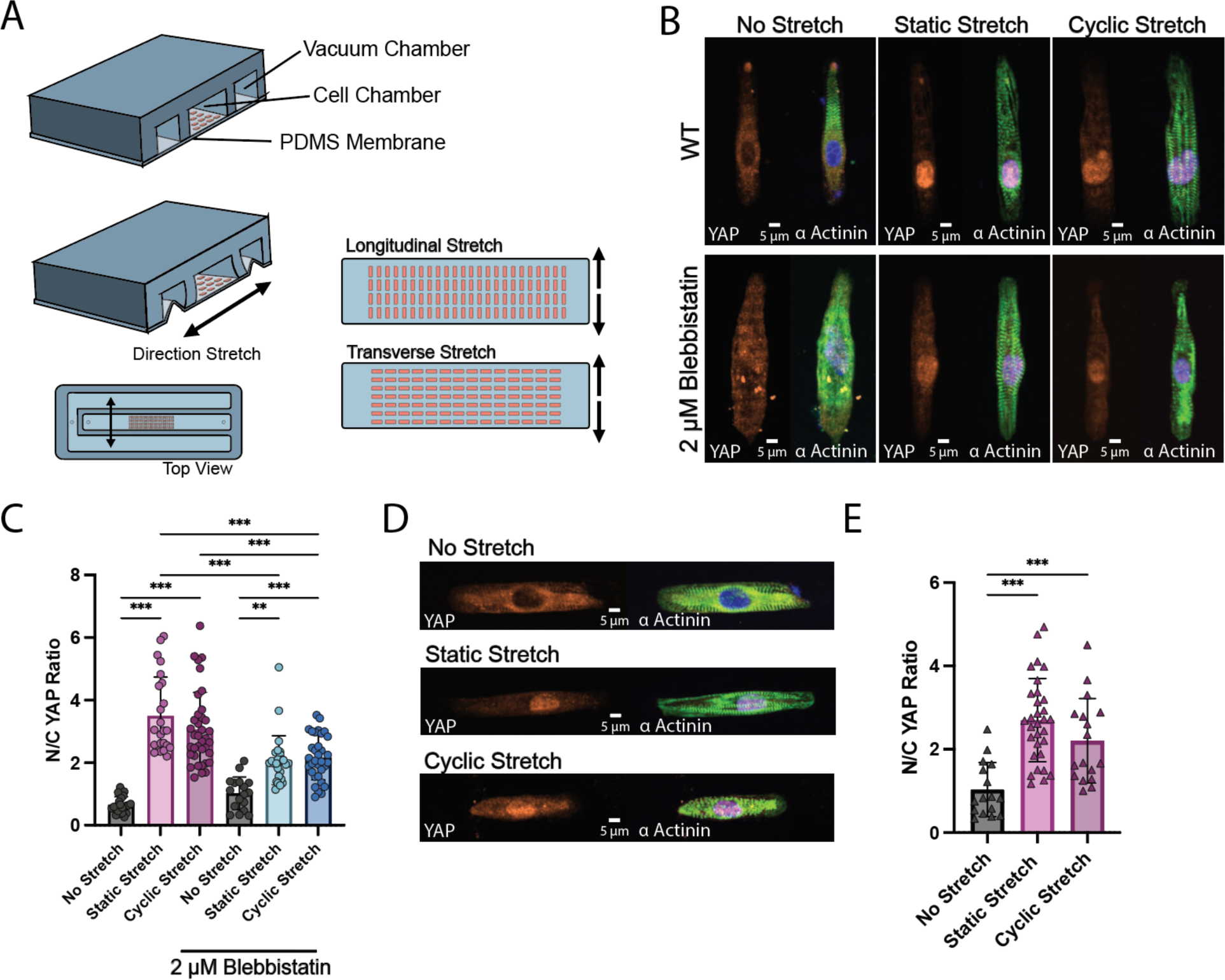
Mechanical deformation in response to uniaxial stretch drives nuclear localization of YAP. (A) Illustration of the Uniaxial Stretch Platform. (B) Representative image of fixed and labeled patterned WT hiPSC-CMs with and without longitudinal stretch (1hz, static and cyclic) and blebbistatin treatment. (C) Quantification of YAP activity from longitudinal stretch experiments inferred by Nuclear to Cytoplasmic (N/C) intensity Ratio. Statistical test: Kruskal-Wallis. (D) Representative image of fixed and labeled patterned WT hiPSC-CMs with and without transverse stretch (1hz, static and cyclic) for 10 mins in the transverse direction. (E) Quantification of YAP activity from transverse stretch experiments inferred by Nuclear to Cytoplasmic (N/C) intensity Ratio. Statistical test: One-way ANOVA.

**Supplement 4.**
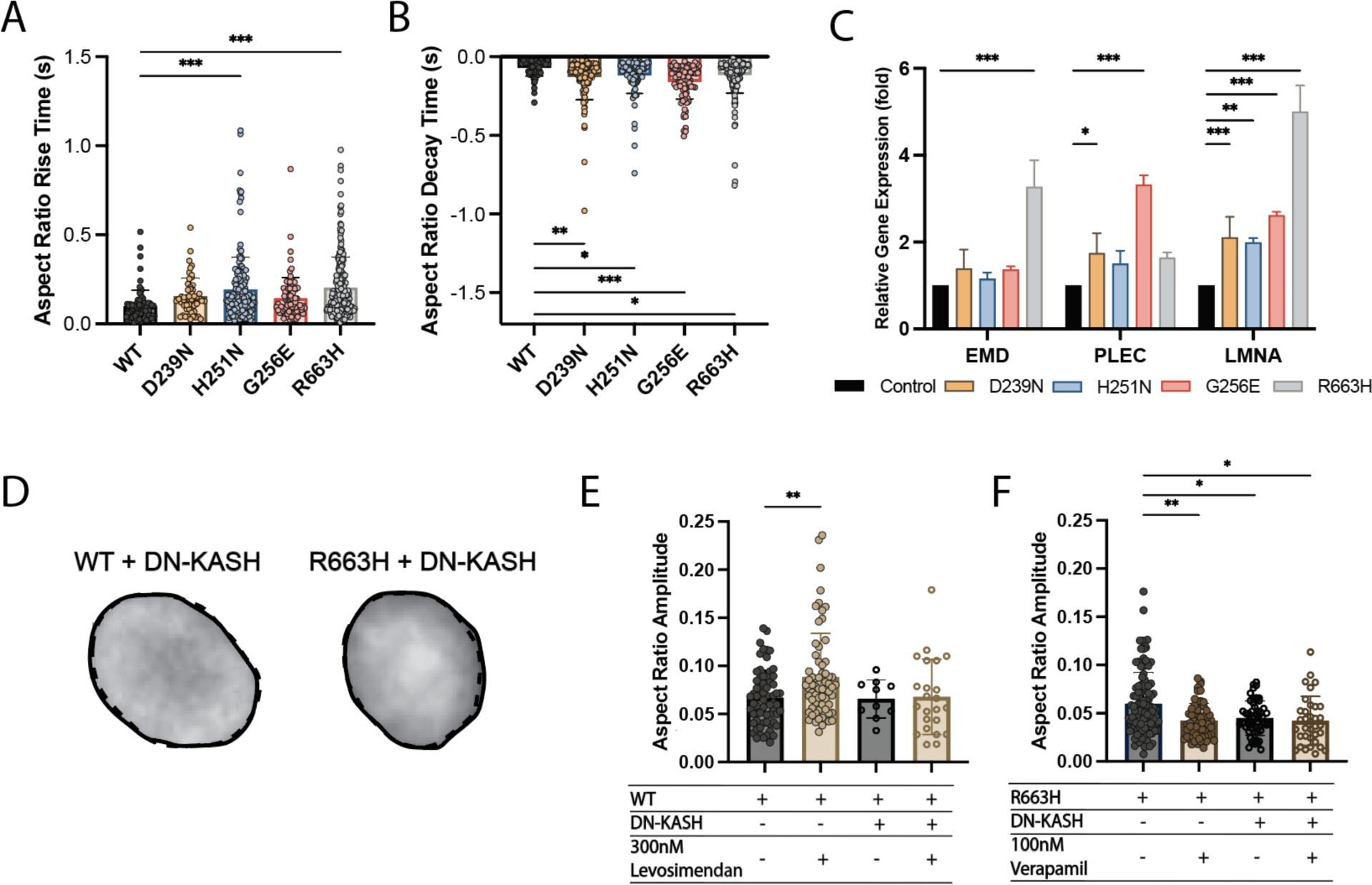
YAP activation in MYH7 mutants requires nuclear mechanical coupling. (A-B) Quantification of nuclear deformation rise time (aspect ratio rise time) and decay time (aspect ratio decay time). Statistical test: Kruskal-Wallis. (C) qPCR results for genes: EMD, PLEC, and LMNA for all four MYH7 mutants (D239N, H251N,G256E and R663H). Statistical test: Kruskal-Wallis. (D) Representative image of WT and MYH7-R663H hiPSC-CM nuclei deforming after DN-KASH treatment (relaxed state (solid,) contracted state (dash)). (E) Quantification of nuclear deformation (aspect ratio amplitude) after DN-KASH and drug treatments for D(60) WT hiPSC-CMs. Statistical test: Kruskal-Wallis. (F) Quantification of nuclear deformation (aspect ratio amplitude) after DN-KASH and drug treatments for MYH7-R663H (right) hiPSC-CMs.

**Supplement 5.**
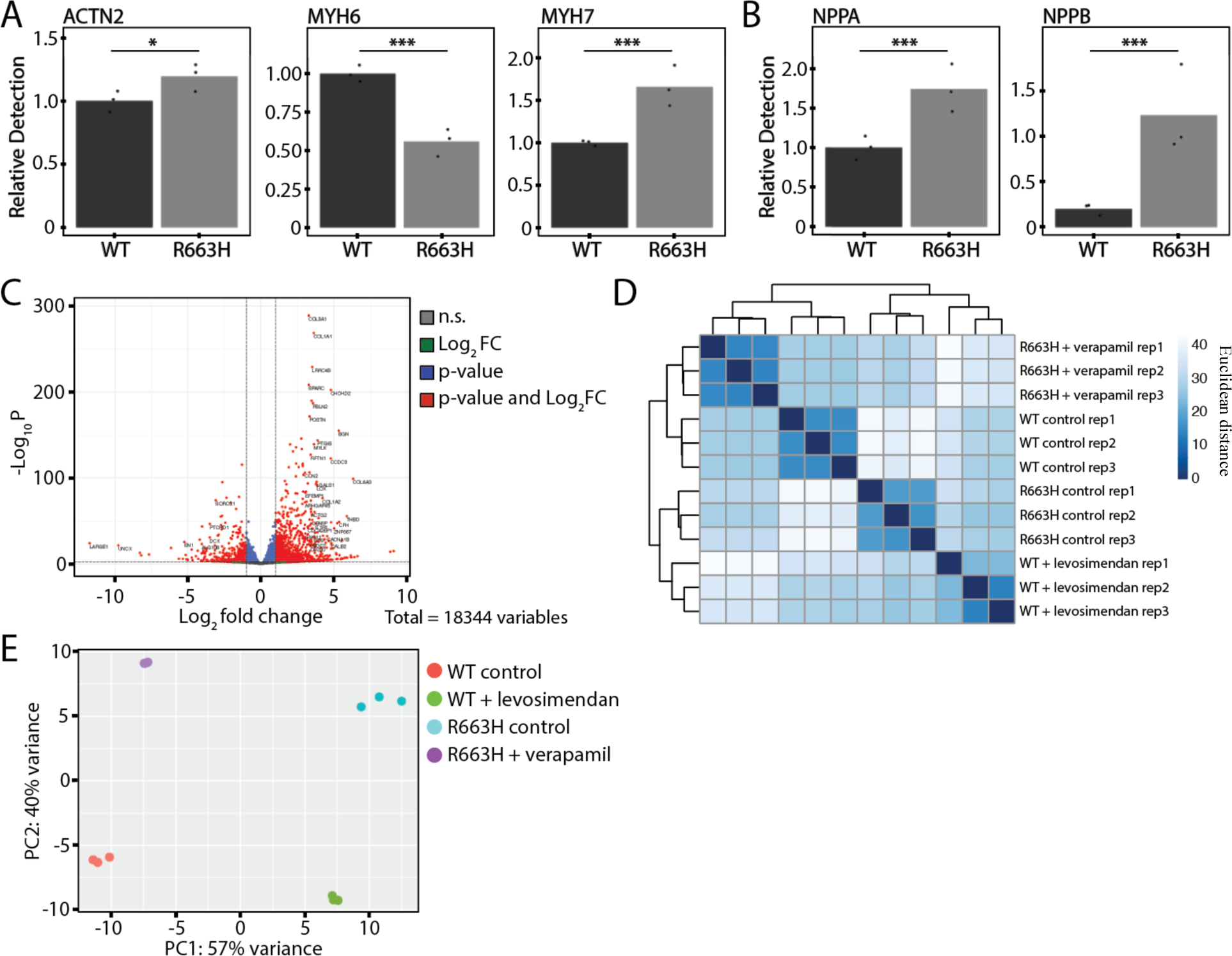
Force modulation alters the transcriptome in WT and MYH7-R663H mutant hiPSC-CMs. (A) RNA-seq results depicting relative detection of known cardiac markers ACTN2, MYH6, and MYH7 for WT and MYH7-R663H hiPSC-CMs. (B) RNA-seq results depicting relative detection of known HCM markers NPPA and NPPB for WT and MYH7-R663H hiPSC-CMs. (C) Volcano plot displaying the top differentially expressed genes (DEGs) for WT and MYH7-R663H hiPSC-CMs. (D) hierarchical clustering heatmap based on Euclidean distance of gene expression from all genes and (E) corresponding PCA plot of WT, WT+levosimendan (300 nM), R663H, and R663H+verapamil (100 nM).

**Supplement 6.**
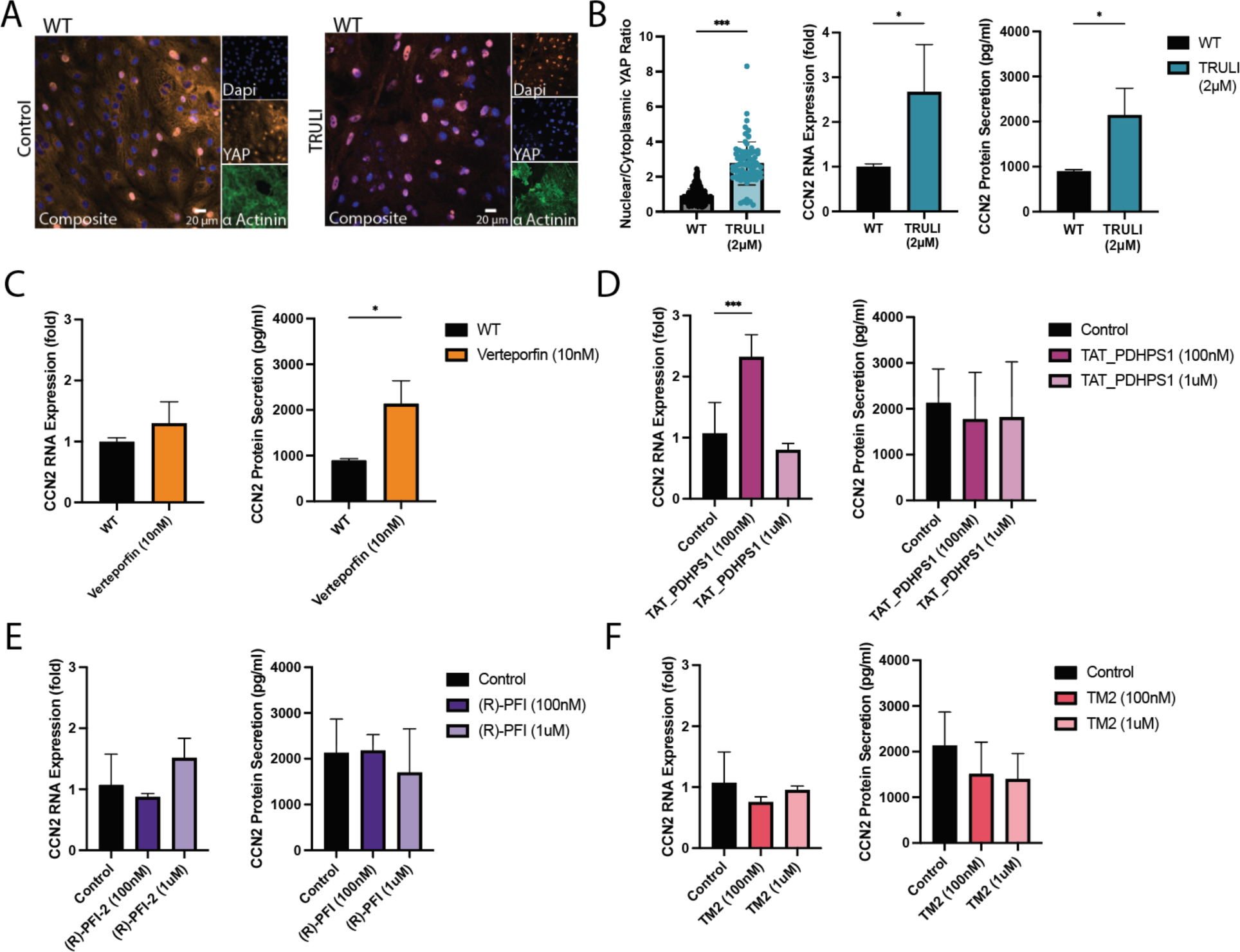
YAP Modulation by Small Molecule. (A) Representative image of fixed and labeled hiPSC-CMs for YAP (red), Alpha Actinin (green) and nuclei (blue) after 24 hr treatment with Hippo Pathway inhibitor TRULI (2 μM, causing the nuclear entry and activation of YAP) examined by immunocytochemistry. (B) (left) Quantification of nuclear to cytoplasmic YAP expression (n= 100 cells for each condition). Statistical test: Mann-Whitney, (middle) qPCR results for CCN2 with and without TRULI treatment (24 hr). Statistical test: Mann-Whitney, (right) Bar plot illustrating CCN2 Elisa results for WT hiPSC-CMs with and without TRULI Treatment (24 hr). Statistical test: unpaired t-test. Data are presented as mean ± STDEV. (C-F) unsuccessful YAP inactivation with small molecules verteporfin, TAT_PDHPS1, (R)-PFI-2, and TM2. Verteporfin was toxic to hiPSC-CMs even at a low dose (10nM). Verteporfin Elisa maybe confounded by cell death. (left) qPCR results for CCN2 with and without drug treatment (24 hr). Statistical test: Mann-Whitney, (right) Bar plot illustrating CCN2 Elisa results for WT hiPSC-CMs with and without drug treatment (24 hr). Statistical test: unpaired t-test. Data are presented as mean ± STDEV.

**Supplement 7.**
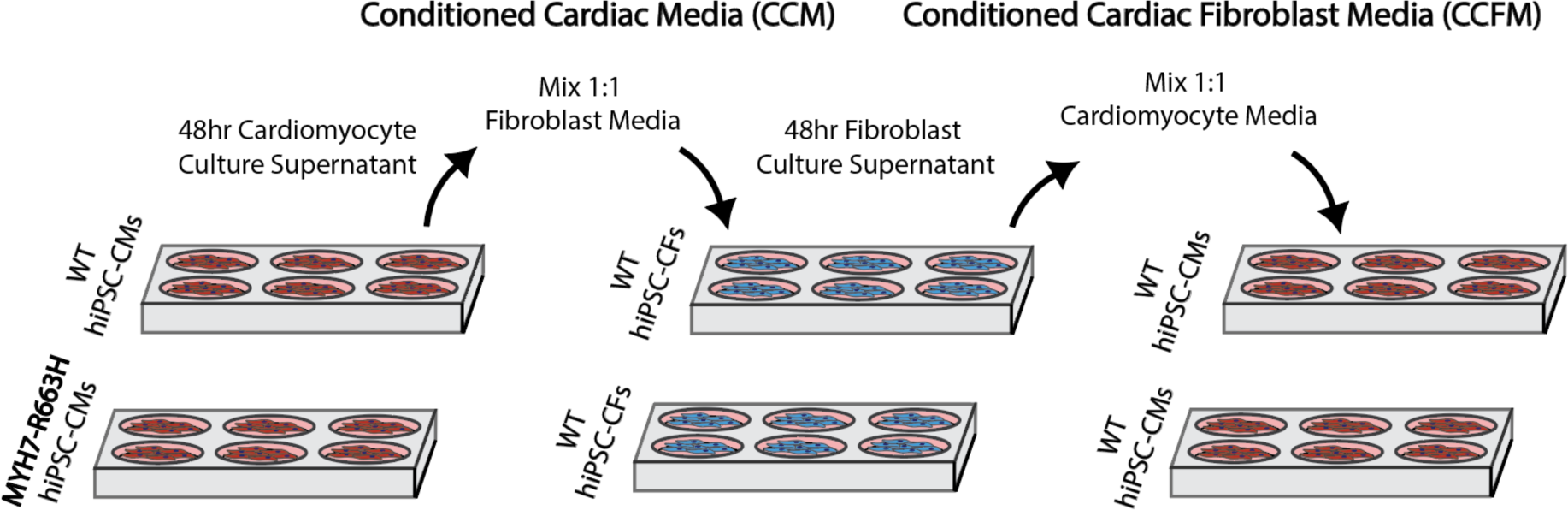
Conditioned Media Experimental Design. (A) Illustration of a reduced order conditioned media experiment to assess the paracrine signaling effects of CCN2 and TGFβ1 on hiPSC-CM size.

**Supplement 8.**
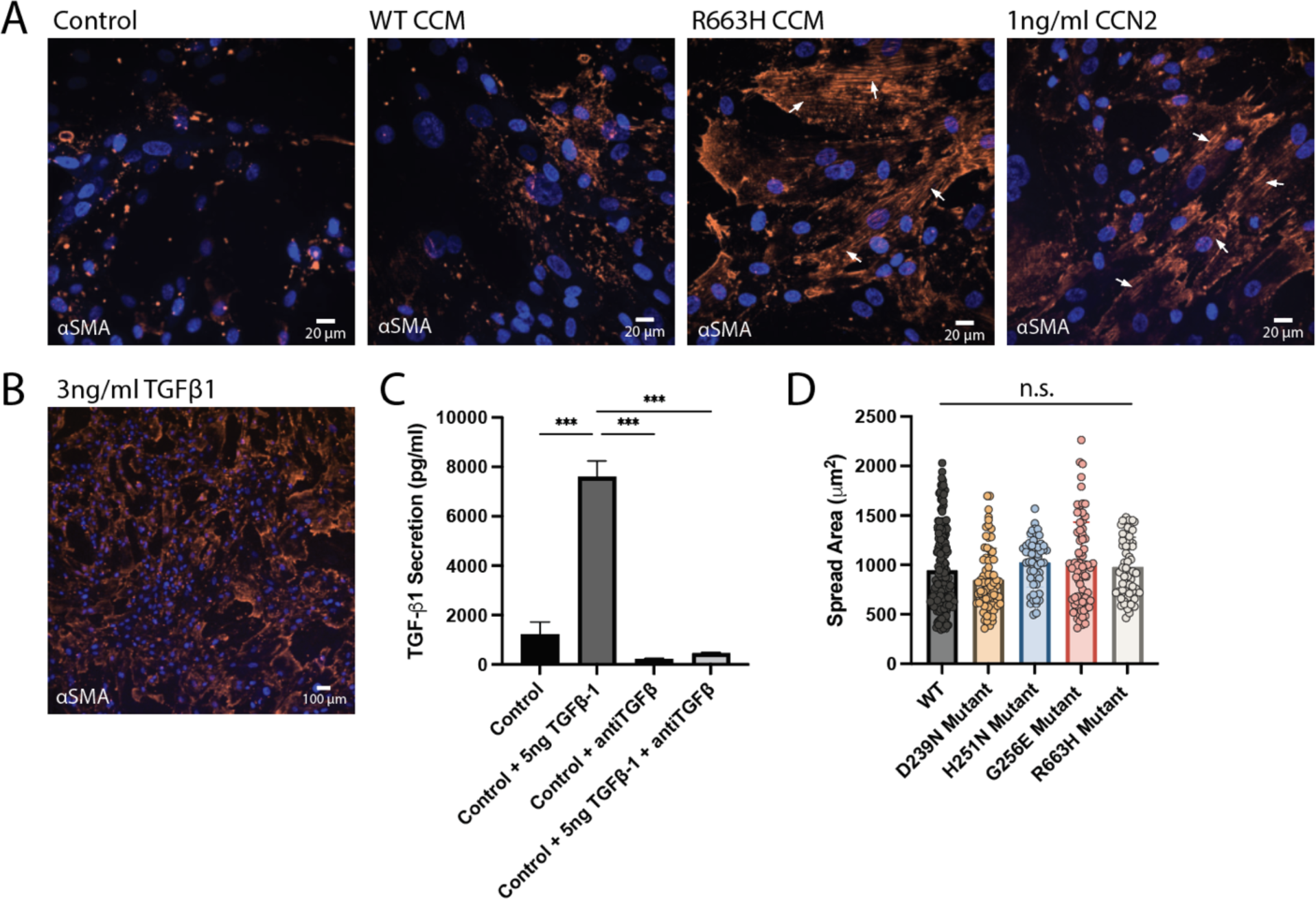
Conditioned media treatment transitions cardiac fibroblasts to myofibroblasts. (A) Representative images of hiPSC-CFs fixed and stained for αSMA, after conditioned media treatment (CCM) or 1 ng/ml CCN2 treatment (144hrs total, 48hr media exchanges). Arrows indicate the incorporation of αSMA into stress fibers, a characteristic of myofibroblast. (B) Representative image of hiPSC-CFs fixed and stained for αSMA, after 3 ng/ml TGFβ1 treatment (144hrs total, 48hr media exchanges). (C) TGFβ1 Elisa results demonstrating appropriate blocking of TGFβ1 with the addition of anti TGFβ1. Statistical test: one-way ANOVA. (D) Quantification of spread area of purified (glucose starved) hiPSC-CM cultures for all four mutations (D239N, H251N, G256E, and R663H) compared to WT.

## Bibliography

1. Ho, C. Y. et al. Genotype and lifetime burden of disease in hypertrophic cardiomyopathy insights from the sarcomeric human cardiomyopathy registry (SHaRe). Circulation 138, 1387–1398 (2018).

2. Harvey, P. A. & Leinwand, L. A. Cellular mechanisms of cardiomyopathy. Journal of Cell Biology vol. 194 355–365 (2011).

3. Semsarian, C., Ingles, J., Maron, M. S. & Maron, B. J. New Perspectives on the Prevalence of Hypertrophic Cardiomyopathy. J. Am. Coll. Cardiol. 65, 1249–1254 (2015).

4. Maron, B. J. & Maron, M. S. Hypertrophic cardiomyopathy. The Lancet vol. 381 242–255 (2013).

5. Marian, A. J. & Braunwald, E. Hypertrophic cardiomyopathy: Genetics, pathogenesis, clinical manifestations, diagnosis, and therapy. Circ. Res. 121, 749–770 (2017).

6. Teekakirikul, P. et al. Cardiac fibrosis in mice with hypertrophic cardiomyopathy is mediated by non-myocyte proliferation and requires Tgf-β. J. Clin. Invest. 120, 3520–3529 (2010).

7. Becker, R. C., Owens, A. P. & Sadayappan, S. Tissue-level inflammation and ventricular remodeling in hypertrophic cardiomyopathy. Journal of Thrombosis and Thrombolysis vol. 49 177–183 (2020).

8. Spudich, J. A. Three perspectives on the molecular basis of hypercontractility caused by hypertrophic cardiomyopathy mutations. Pflugers Archiv European Journal of Physiology vol. 471 701–717 (2019).

9. Spudich, J. A. Hypertrophic and dilated cardiomyopathy: four decades of basic research on muscle lead to potential therapeutic approaches to these devastating genetic diseases. Biophys. J. 106, 1236–49 (2014).

10. Xu, Q., Dewey, S., Nguyen, S. & Gomes, A. V. Malignant and benign mutations in familial cardiomyopathies: insights into mutations linked to complex cardiovascular phenotypes. J. Mol. Cell. Cardiol. 48, 899–909 (2010).

11. Lam, C. K. et al. Identifying the transcriptome signatures of calcium channel blockers in human induced pluripotent stem cell-derived cardiomyocytes. Circ. Res. 125, 212–222 (2019).

12. Gersh, B. J. et al. 2011 ACCF/AHA guideline for the diagnosis and treatment of hypertrophic cardiomyopathy: Executive summary: A report of the American College of cardiology foundation/American heart association task force on practice guidelines. Circulation 124, 2761–2796 (2011).

13. Aolo, P. et al. The Management of Hypertrophic Cardiomyopathy. https://doi.org/10.1056/NEJM199703133361107 44, 195–202 (1997).

14. Wu, H. et al. Modelling diastolic dysfunction in induced pluripotent stem cell-derived cardiomyocytes from hypertrophic cardiomyopathy patients. European Heart Journal vol. 40 3685–3695 (Oxford University Press, 2019).

15. Olivotto, I. et al. Mavacamten for treatment of symptomatic obstructive hypertrophic cardiomyopathy (EXPLORER-HCM): a randomised, double-blind, placebo-controlled, phase 3 trial. Lancet 396, 759–769 (2020).

16. Toepfer, C. N. et al. Myosin Sequestration Regulates Sarcomere Function, Cardiomyocyte Energetics, and Metabolism, Informing the Pathogenesis of Hypertrophic Cardiomyopathy. Circulation 141, 828 (2020).

17. Cho, S. et al. Mechanosensing by the Lamina Protects against Nuclear Rupture, DNA Damage, and Cell-Cycle Arrest. Dev. Cell 49, 920–935.e5 (2019).

18. Ghosh, S. et al. Deformation Microscopy for Dynamic Intracellular and Intranuclear Mapping of Mechanics with High Spatiotemporal Resolution. Cell Rep. 27, 1607–1620.e4 (2019).

19. Münch, J. & Abdelilah-Seyfried, S. Sensing and Responding of Cardiomyocytes to Changes of Tissue Stiffness in the Diseased Heart. Front. cell Dev. Biol. 9, (2021).

20. Engler, A. J., Sen, S., Sweeney, H. L. & Discher, D. E. Matrix Elasticity Directs Stem Cell Lineage Specification. Cell 126, 677–689 (2006).

21. De Belly, H., Paluch, E. K. & Chalut, K. J. Interplay between mechanics and signalling in regulating cell fate. Nat. Rev. Mol. Cell Biol. 2022 237 23, 465–480 (2022).

22. Wang, N., Tytell, J. D. & Ingber, D. E. Mechanotransduction at a distance: mechanically coupling the extracellular matrix with the nucleus. Nat. Rev. Mol. Cell Biol. 2009 101 10, 75–82 (2009).

23. Martino, F., Perestrelo, A. R., Vinarský, V., Pagliari, S. & Forte, G. Cellular Mechanotransduction: From Tension to Function. Front. Physiol. 9, (2018).

24. Elosegui-Artola, A. et al. Force Triggers YAP Nuclear Entry by Regulating Transport across Nuclear Pores. Cell 171, 1397–1410.e14 (2017).

25. Song, Y. et al. Transient nuclear deformation primes epigenetic state and promotes cell reprogramming. Nat. Mater. 2022 2110 21, 1191–1199 (2022).

26. Seelbinder, B., et al. The Nucleus Mediates Mechanosensitive Reorganization of Epigenetically Marked Chromatin During Cardiac Maturation and Pathology. bioRxiv 455600 (2018) doi:10.1101/455600.

27. Ma, S., Meng, Z., Chen, R. & Guan, K.-L. The Hippo Pathway: Biology and Pathophysiology. Annu. Rev. Biochem. 88, 577–604 (2019).

28. Zhao, B. et al. TEAD mediates YAP-dependent gene induction and growth control. Genes Dev. 22, 1962–1971 (2008).

29. Meng, Z., Moroishi, T. & Guan, K. L. Mechanisms of Hippo pathway regulation. Genes and Development vol. 30 1–17 (2016).

30. Hill, M. C. et al. Integrated multi-omic characterization of congenital heart disease. Nat. 2022 6087921 608, 181–191 (2022).

31. Wang, P. et al. The alteration of Hippo/YAP signaling in the development of hypertrophic cardiomyopathy. Basic Res. Cardiol. 109, 1–11 (2014).

32. Rosenkranz, S. et al. Alterations of β-adrenergic signaling and cardiac hypertrophy in transgenic mice overexpressing TGF-β1. Am. J. Physiol. - Hear. Circ. Physiol. 283, (2002).

33. Dobaczewski, M., Chen, W. & Frangogiannis, N. G. Transforming Growth Factor (TGF)-β signaling in cardiac remodeling. J. Mol. Cell. Cardiol. 51, 600 (2011).

34. Ma, Y. et al. Transforming growth factor β: A potential biomarker and therapeutic target of ventricular remodeling. Oncotarget 8, 53780 (2017).

35. Plouffe, S. W., Hong, A. W. & Guan, K. L. Disease implications of the Hippo/YAP pathway. Trends in Molecular Medicine vol. 21 212–222 (2015).

36. Zanconato, F., Cordenonsi, M. & Piccolo, S. YAP/TAZ at the Roots of Cancer. Cancer Cell vol. 29 783–803 (2016).

37. Zhao, B., Li, L., Lei, Q. & Guan, K. L. The Hippo-YAP pathway in organ size control and tumorigenesis: An updated version. Genes and Development vol. 24 862–874 (2010).

38. Kim, C.-L., Choi, S.-H. & Mo, J.-S. Role of the Hippo Pathway in Fibrosis and Cancer. Cells 8, 468 (2019).

39. Roest, A. S. Vander et al. Hypertrophic cardiomyopathy β-cardiac myosin mutation (P710R) leads to hypercontractility by disrupting super relaxed state. Proc. Natl. Acad. Sci. U. S. A. 118, e2025030118 (2021).

40. Garg, P. et al. Genome Editing of Induced Pluripotent Stem Cells to Decipher Cardiac Channelopathy Variant. J. Am. Coll. Cardiol. 72, 62–75 (2018).

41. Li, W. et al. Disease Phenotypes and Mechanisms of iPSC-Derived Cardiomyocytes From Brugada Syndrome Patients With a Loss-of-Function SCN5A Mutation. Front. Cell Dev. Biol. 8, 1181 (2020).

42. Park, S. J. et al. Insights Into the Pathogenesis of Catecholaminergic Polymorphic Ventricular Tachycardia From Engineered Human Heart Tissue. Circulation 140, 390–404 (2019).

43. Sun, N. et al. Patient-Specific Induced Pluripotent Stem Cell as a Model for Familial Dilated Cardiomyopathy. Sci. Transl. Med. 4, 130ra47 (2012).

44. Lan, F. et al. Abnormal calcium handling properties underlie familial hypertrophic cardiomyopathy pathology in patient-specific induced pluripotent stem cells. Cell Stem Cell 12, 101–113 (2013).

45. Li, J., Feng, X. & Wei, X. Modeling hypertrophic cardiomyopathy with human cardiomyocytes derived from induced pluripotent stem cells. Stem Cell Res. Ther. 13, (2022).

46. Gruver, E. J. et al. Familial hypertrophic cardiomyopathy and atrial fibrillation caused by Arg663His beta-cardiac myosin heavy chain mutation. Am. J. Cardiol. 83, 13–18 (1999).

47. Fananapazir, L., Dalakas, M. C., Cyran, F., Cohn, G. & Epstein, N. D. Missense mutations in the beta-myosin heavy-chain gene cause central core disease in hypertrophic cardiomyopathy. Proc. Natl. Acad. Sci. U. S. A. 90, 3993–3997 (1993).

48. Adhikari, A. S. et al. Early-Onset Hypertrophic Cardiomyopathy Mutations Significantly Increase the Velocity, Force, and Actin-Activated ATPase Activity of Human β-Cardiac Myosin. Cell Rep. 17, 2857–2864 (2016).

49. Kaski, J. P. et al. Prevalence of sarcomere protein gene mutations in preadolescent children with hypertrophic cardiomyopathy. Circ. Cardiovasc. Genet. 2, 436–441 (2009).

50. Sarkar, S. S. et al. The hypertrophic cardiomyopathy mutations R403Q and R663H increase the number of myosin heads available to interact with actin. Sci. Adv. 6, (2020).

51. Lian, X. et al. Robust cardiomyocyte differentiation from human pluripotent stem cells via temporal modulation of canonical Wnt signaling. Proc. Natl. Acad. Sci. U. S. A. 109, E1848–E1857 (2012).

52. Monroe, T. O. et al. YAP Partially Reprograms Chromatin Accessibility to Directly Induce Adult Cardiogenesis In Vivo. Dev. Cell 48, 765–779.e7 (2019).

53. Yang, Y. et al. MIR-206 Mediates YAP-Induced Cardiac Hypertrophy and Survival. Circ. Res. 117, 891–904 (2015).

54. Dias, T. P. et al. Biophysical study of human induced Pluripotent Stem Cell-Derived cardiomyocyte structural maturation during long-term culture. Biochem. Biophys. Res. Commun. 499, 611–617 (2018).

55. Zhao, B. et al. Inactivation of YAP oncoprotein by the Hippo pathway is involved in cell contact inhibition and tissue growth control. Genes Dev. 21, 2747–2761 (2007).

56. Guo, Q. et al. The WW domains dictate isoform-specific regulation of YAP1 stability and pancreatic cancer cell malignancy. Theranostics 10, 4422–4436 (2020).

57. Ribeiro, A. J. S. S. et al. Contractility of single cardiomyocytes differentiated from pluripotent stem cells depends on physiological shape and substrate stiffness. Proc. Natl. Acad. Sci. 112, 12705–12710 (2015).

58. Ribeiro, A. J. S. et al. Multi-Imaging Method to Assay the Contractile Mechanical Output of Micropatterned Human iPSC-Derived Cardiac Myocytes. Circ Res 120, 1572–1583 (2017).

59. Muñoz, J. J. A. M. et al. Time-regulated transcripts with the potential to modulate human pluripotent stem cell-derived cardiomyocyte differentiation. Stem Cell Res. Ther. 13, 1–27 (2022).

60. Wang, L. et al. Hypertrophic cardiomyopathy-linked mutation in troponin T causes myofibrillar disarray and pro-arrhythmic action potential changes in human iPSC cardiomyocytes. J. Mol. Cell. Cardiol. 114, 320–327 (2018).

61. Kawana, M., Sarkar, S. S., Sutton, S., Ruppel, K. M. & Spudich, J. A. Biophysical properties of human b-cardiac myosin with converter mutations that cause hypertrophic cardiomyopathy. Sci. Adv. 3, e1601959 (2017).

62. Trivedi, D. V., Adhikari, A. S., Sarkar, S. S., Ruppel, K. M. & Spudich, J. A. Hypertrophic cardiomyopathy and the myosin mesa: viewing an old disease in a new light. Biophys. Rev. 10, 27–48 (2018).

63. Ribeiro, M. C. et al. A cardiomyocyte show of force: A fluorescent alpha-actinin reporter line sheds light on human cardiomyocyte contractility versus substrate stiffness. J. Mol. Cell. Cardiol. 141, 54–64 (2020).

64. Haikala, H., Levijoki, J. & Lindén, I. B. Troponin C-mediated calcium sensitization by levosimendan accelerates the proportional development of isometric tension. J. Mol. Cell. Cardiol. 27, 2155–2165 (1995).

65. Lam, C. K. et al. Identifying the transcriptome signatures of calcium channel blockers in human induced pluripotent stem cell-derived cardiomyocytes. Circ. Res. 125, 212–222 (2019).

66. Hart, K. C. et al. An Easy-to-Fabricate Cell Stretcher Reveals Density-Dependent Mechanical Regulation of Collective Cell Movements in Epithelia. Cell. Mol. Bioeng. 14, 569–581 (2021).

67. Koushki, N. et al. Lamin A redistribution mediated by nuclear deformation determines dynamic localization of YAP. bioRxiv 2020.03.19.998708 (2020) doi:10.1101/2020.03.19.998708.

68. Gaspar, P. & Tapon, N. Sensing the local environment: Actin architecture and Hippo signalling. Current Opinion in Cell Biology vol. 31 74–83 (2014).

69. Vite, A., Zhang, C., Yi, R., Emms, S. & Radice, G. L. α-catenin-dependent cytoskeletal tension controls yap activity in the heart. Dev. 145, (2018).

70. Piccolo, S., Dupont, S. & Cordenonsi, M. The Biology of YAP/TAZ: Hippo Signaling and Beyond. Physiol. Rev. 94, 1287–1312 (2014).

71. Cho, S., Irianto, J. & Discher, D. E. Mechanosensing by the nucleus: From pathways to scaling relationships. J. Cell Biol. 216, 305 (2017).

72. Kanehisa, M., Furumichi, M., Tanabe, M., Sato, Y. & Morishima, K. KEGG: New perspectives on genomes, pathways, diseases and drugs. Nucleic Acids Res. 45, D353–D361 (2017).

73. Kim, N. G. & Gumbiner, B. M. Adhesion to fibronectin regulates Hippo signaling via the FAK-Src-PI3K pathway. J. Cell Biol. 210, 503–515 (2015).

74. Dorn, L. E., Petrosino, J. M., Wright, P. & Accornero, F. CTGF/CCN2 is an autocrine regulator of cardiac fibrosis. J. Mol. Cell. Cardiol. 121, 205 (2018).

75. Vainio, L. E. et al. Connective Tissue Growth Factor Inhibition Enhances Cardiac Repair and Limits Fibrosis After Myocardial Infarction. JACC Basic to Transl. Sci. 4, 83–94 (2019).

76. Lipson, K. E., Wong, C., Teng, Y. & Spong, S. CTGF is a central mediator of tissue remodeling and fibrosis and its inhibition can reverse the process of fibrosis. Fibrogenesis Tissue Repair 5, 1–8 (2012).

77. Rupérez, M. et al. Connective tissue growth factor is a mediator of angiotensin II-induced fibrosis. Circulation 108, 1499–1505 (2003).

78. Leask, A. Et tu, CCN1…. J. Cell Commun. Signal. 14, 355–356 (2020).

79. Qin, Z., He, T., Guo, C. & Quan, T. Age-Related Downregulation of CCN2 Is Regulated by Cell Size in a YAP/TAZ-Dependent Manner in Human Dermal Fibroblasts: Impact on Dermal Aging. JID Innov. Ski. Sci. from Mol. to Popul. Heal. 2, 100111 (2022).

80. Kastan, N. et al. Small-molecule inhibition of Lats kinases may promote Yap-dependent proliferation in postmitotic mammalian tissues. Nat. Commun. 12, 3100 (2021).

81. Giraud, J. et al. Verteporfin targeting YAP1/TAZ-TEAD transcriptional activity inhibits the tumorigenic properties of gastric cancer stem cells. Int. J. cancer 146, 2255–2267 (2020).

82. Hu, L. et al. Discovery of a new class of reversible TEA domain transcription factor inhibitors with a novel binding mode. Elife 11, (2022).

83. Pan, X. et al. Peptide PDHPS1 Inhibits Ovarian Cancer Growth through Disrupting YAP Signaling. Mol. Cancer Ther. 21, 1160–1170 (2022).

84. Barsyte-Lovejoy, D. et al. (R)-PFI-2 is a potent and selective inhibitor of SETD7 methyltransferase activity in cells. Proc. Natl. Acad. Sci. U. S. A. 111, 12853–12858 (2014).

85. Dorn, L. E., Petrosino, J. M., Wright, P. & Accornero, F. CTGF/CCN2 is an autocrine regulator of cardiac fibrosis. J. Mol. Cell. Cardiol. 121, 205–211 (2018).

86. Meng, Q. et al. Myofibroblast-Specific TGFβ Receptor II Signaling in the Fibrotic Response to Cardiac Myosin Binding Protein C-Induced Cardiomyopathy. Circ. Res. 123, 1285–1297 (2018).

87. Hinz, B. Formation and function of the myofibroblast during tissue repair. J. Invest. Dermatol. 127, 526–537 (2007).

88. Prunotto, M. et al. Stable incorporation of α-smooth muscle actin into stress fibers is dependent on specific tropomyosin isoforms. Cytoskeleton (Hoboken). 72, 257–267 (2015).

89. Li, J. M. & Brooks, G. Differential protein expression and subcellular distribution of TGFbeta1, beta2 and beta3 in cardiomyocytes during pressure overload-induced hypertrophy. J. Mol. Cell. Cardiol. 29, 2213–2224 (1997).

90. Dobaczewski, M., Chen, W. & Frangogiannis, N. G. Transforming growth factor (TGF)-β signaling in cardiac remodeling. J. Mol. Cell. Cardiol. 51, 600–606 (2011).

91. Serini, G. et al. The fibronectin domain ED-A is crucial for myofibroblastic phenotype induction by transforming growth factor-beta1. J. Cell Biol. 142, 873–881 (1998).

92. Rosenkranz, S. et al. Alterations of beta-adrenergic signaling and cardiac hypertrophy in transgenic mice overexpressing TGF-beta(1). Am. J. Physiol. Heart Circ. Physiol. 283, (2002).

93. Nakano, H. et al. Glucose inhibits cardiac muscle maturation through nucleotide biosynthesis. Elife 6, 1–23 (2017).

94. Ni, X. et al. Single-cell analysis reveals the purification and maturation effects of glucose starvation in hiPSC-CMs. Biochem. Biophys. Res. Commun. 534, 367–373 (2021).

95. Kreitzer, F. R. et al. A robust method to derive functional neural crest cells from human pluripotent stem cells. Am. J. Stem Cells 2, 119–131 (2013).

96. Lan, F. et al. Abnormal Calcium Handling Properties Underlie Familial Hypertrophic Cardiomyopathy Pathology in Patient-Specific Induced Pluripotent Stem Cells. Cell Stem Cell 12, 101–113 (2013).

97. Sharma, A. et al. Derivation of Highly Purified Cardiomyocytes from Human Induced Pluripotent Stem Cells Using Small Molecule-modulated Differentiation and Subsequent Glucose Starvation. J. Vis. Exp. 2015, 52628 (2015).

98. Whitehead, A. J., Hocker, J. D., Ren, B. & Engler, A. J. Improved epicardial cardiac fibroblast generation from iPSCs. J. Mol. Cell. Cardiol. 164, 58–68 (2022).

99. Ribeiro, A. J. S. et al. Multi-Imaging Method to Assay the Contractile Mechanical Output of Micropatterned Human iPSC-Derived Cardiac Myocytes. Circ. Res. 120, 1572–1583 (2017).

100. Love, M. I., Huber, W. & Anders, S. Moderated estimation of fold change and dispersion for RNA-seq data with DESeq2. Genome Biol. 15, (2014).

101. Stephens, M. False discovery rates: a new deal. Biostatistics 18, 275 (2017).

102. Ritchie, M. E. et al. limma powers differential expression analyses for RNA-sequencing and microarray studies. Nucleic Acids Res. 43, e47 (2015).

103. Sherman, B. T. et al. DAVID: a web server for functional enrichment analysis and functional annotation of gene lists (2021 update). Nucleic Acids Res. 50, W216–W221 (2022).

